# The limitations of non-mechanistic methods for characterizing pathogen-pathogen interactions: A simulation study

**DOI:** 10.64898/2025.12.22.695879

**Authors:** Sarah C. Kramer, Sarah Pirikahu, Cana Kussmaul, Lulla Opatowski, Matthieu Domenech de Cellès

## Abstract

Pathogen-pathogen interactions occur when infection with one pathogen influences one’s chance of infection or disease due to another. Increasingly, evidence suggests that interactions are a common feature of infectious disease epidemiology. However, due to both the nonlinearities and stochasticity inherent to infectious disease transmission, and the frequency of confounding (e.g., by shared seasonal forcing), simple, purely statistical methods for characterizing interactions may be prone to failure. Here, we perform a simulation study to evaluate several more complex non-mechanistic approaches for inferring causality from time series data: generalized additive models (GAMs), Granger causality, transfer entropy, and convergent cross-mapping (CCM). Specifically, we use a two-pathogen mechanistic transmission model, calibrated to produce dynamics resembling outbreaks of influenza and respiratory syncytial virus (RSV), to generate synthetic datasets with a range of values for interaction strength and duration. We then apply each method to all synthetic datasets. We find that Granger causality, transfer entropy, and CCM all fail to consistently infer whether data contain signal of an interaction; in particular, methods tend to incorrectly identify interactions where none are modeled (average sensitivity = 80.6%, 92.1%, 72.1%, respectively; average specificity = 31.0%, 33.3%, 33.1%). Furthermore, we find little to no association between point estimates from each method and true interaction strength. In contrast, GAMs infer the existence of interactions more accurately than the other methods (sensitivity = 85.2%, specificity = 72.5%), and consistently yield larger point estimates for stronger interactions. However, their practical utility is limited by an inability to evaluate interaction asymmetry (i.e., whether the effect of pathogen A on pathogen B is identical to that of B on A). Overall performance patterns were similar when methods were applied to two real-world datasets from Hong Kong and Canada. We conclude that accurately and comprehensively characterizing pathogen-pathogen interactions based on outbreak data remains a significant challenge. For this reason, it is critical that any proposed methods be rigorously evaluated before being used to draw conclusions about interactions.

**Author Summary:** Pathogen-pathogen interactions occur when infection with one pathogen either increases or decreases a person’s risk of infection or illness due to a second, distinct pathogen. Because interactions affect several common human pathogens, including influenza and SARS-CoV-2, a better understanding of interactions could improve epidemic control. However, past work has shown that simple methods commonly used to study interactions can lead to inaccurate conclusions. Here, we tested four methods frequently used in other fields, including ecology and neuroscience, to see whether they may also be useful for identifying interactions. Specifically, we tested each method using simulated outbreak data generated from a mathematical model. We found that most methods struggled to correctly determine whether an interaction effect was present; in particular, methods often falsely identified interactions when none occurred. Although one of the tested methods, generalized additive models, performed comparatively well at identifying interactions, it provided relatively little additional information about the interactions. Because pathogen-pathogen interactions are so challenging to study, it is important that researchers rigorously test methods before applying them to interactions, so as not to publish potentially misleading results. More broadly, a complete understanding of interactions will likely require a variety of approaches, including both laboratory and modeling studies.

## Introduction

Although infectious diseases are most commonly studied in isolation, pathogen-pathogen interactions, in which infection with one pathogen modifies the risk of infection or disease due to another pathogen, are rapidly gaining more attention. Throughout the literature, the term “interaction” is applied broadly and sometimes inconsistently: population-level phenomena (e.g., reductions in cases of subsequent pathogens due to people remaining at home during convalescence) and even purely statistical associations are sometimes included. In this work, we use the term to refer exclusively to interactions arising due to biological, within-host mechanisms. These biological mechanisms are varied, and include both direct (e.g., competition for resources) and indirect (e.g., modulation of host cells or the immune response) processes (1,2). Recent evidence suggests that a complex web of such interactions exists between respiratory viruses, including influenza virus, respiratory syncytial virus (RSV), rhinovirus, human metapneumovirus (HMPV), and severe acute respiratory syndrome coronavirus 2 (SARS-CoV-2) (1,3–12). Many of these viruses may in turn affect infection with bacteria like *S. pneumoniae* and *Haemophilus influenzae* (13,14), while the impact of immunosuppression induced by viruses like HIV and measles on susceptibility to a wide range of pathogens has long been recognized (15,16). Indeed, it seems likely that interactions are the norm, rather than the exception, in infectious disease epidemiology. Although interactions as defined here occur at the level of the individual, their effects on outbreak dynamics at the population level can be substantial (1,2). For this reason, characterizing these interactions is a necessary step in improving our understanding of the epidemiology of a wide range of infectious diseases.

Unfortunately, pathogen-pathogen interactions are difficult to characterize. In part, this is because interactions themselves are complex: they can vary in strength and duration, may be positive or negative (i.e., causing either an increase or a decrease in susceptibility to or severity of infection with a subsequent pathogen), and may be symmetric (i.e., having an equivalent impact in both directions) or asymmetric (1). To fully understand an interaction, each of these components must be described. A further challenge is introduced when attempting to study interactions using surveillance data. Because of nonlinearities and stochasticity in the underlying transmission dynamics of the pathogens under study, statistical patterns in the observed number of cases or deaths do not necessarily reflect patterns in the underlying drivers (17,18). Additionally, confounders, such as shared seasonal forcing, may create associations between observed time series that are not due to interaction effects. For these reasons, simple statistical associations identified using observed data are rarely informative about the presence or nature of an underlying, biological interaction: measures of coinfection prevalence systematically underestimate the true strength of an interaction (19), while phase differences are not a reliable indicator of whether an interaction is positive or negative (20). Despite these findings, these simple approaches remain common in the interactions literature. Although fitting mechanistic transmission models to empirical data is a promising alternative (1–3), these methods are computationally intensive, and the development of appropriate models requires a good understanding of the natural history and transmission dynamics of the included pathogens.

A variety of non-mechanistic methods have been designed to infer causality between time series, including Granger causality (21,22), transfer entropy (23), and, most recently, convergent cross-mapping (CCM) (24). Meanwhile, methods like generalized additive models (GAMs) (25) can flexibly account for potential confounders while simultaneously inferring relationships between variables. Compared to a mechanistic modeling approach, these methods are relatively easy to implement, and require much less understanding of the underlying transmission dynamics. These methods are popular in many fields, such as econometrics (26,27), neuroscience (28,29) and ecology and environmental science (24,30–33). They have also been used in infectious disease epidemiology, specifically in elucidating the impact of climatic drivers (18,34) and transmission dynamics by age (35). However, very few studies have evaluated them in the context of pathogen-pathogen interactions. Cobey and Baskerville (36) evaluated CCM as a tool for identifying lifelong cross-immunity between pathogen strains, but did not consider shorter-term interactions. Meanwhile, Randuineau (37) tested Granger causality and transfer entropy, but focused only on unidirectional interactions.

We note preemptively that the above methods were not developed specifically with pathogen-pathogen interactions in mind, and that observed case data commonly available for epidemiologic research may not always meet every assumption of these methods (see Methods and Discussion for more details). With that said, some of these methods have proven surprisingly effective in scenarios where their assumptions don’t necessarily hold (30). Furthermore, it is unfortunately common in infectious disease epidemiology research to see methods applied in situations where their assumptions are violated. Indeed, Barrero Guevara et al. found that time series regression was commonly used in studies evaluating the influence of weather on infectious disease transmission, despite the fact that associations between weather variables and downstream disease incidence do not necessarily reflect the association between weather and transmission rates (17,38). Similarly, despite the work discussed above showing that prevalence ratios and phase differences are not reliable as indicators of pathogen-pathogen interactions, these methods remain popular. Given that recent research has already attempted to apply some of the methods tested here to pathogen-pathogen interactions (39,40), we consider a formal test of these methods to be of critical importance, even if they may not be perfectly applicable.

In this work, we conducted a simulation study to assess whether several commonly-used non-mechanistic methods can correctly classify the interaction effect between two pathogens when confronted with synthetic data generated by a mathematical model. More specifically, we evaluate the extent to which a method returning a statistically significant result reliably corresponds to a true, biological interaction between the modeled pathogens (and, conversely, the extent to which null results are indicative of a lack of such an interaction). Furthermore, we assessed whether the magnitude of estimates produced by each method could provide information about the relative strength of the underlying interaction. Methods were tested for interactions with a range of strengths and durations, and, where possible, implementations controlling for the shared effect of seasonal forcing on both pathogens were also run. Overall, we aim to determine whether the tested methods show promise in identifying and characterizing interactions based on real-world outbreak data.

## Methods

### Implementation

All analyses were conducted in R version 4.4.0 (41). All code used for this project has been published on GitHub (42).

### Mathematical Model

#### Transmission Model

We generated ten years of synthetic data for two cocirculating pathogens using a compartmental SEITRSxSEITRS model, where S stands for susceptible, E for exposed but not infectious, I for infected and able to transmit, T for a transient period post-infection, and R for recovered and immune. Here, individuals’ infection status relative to both pathogens is tracked simultaneously (e.g., individuals in compartment *X*_*SI*_ are susceptible to pathogen A and infected with pathogen B). The interaction is modeled as a change in the force of infection for pathogen B among those currently infected (I) or recently infected (T) with pathogen A, and vice versa, consistent with evidence suggesting that pathogen-pathogen interactions can arise due to infection with one pathogen influencing susceptibility to another (4,5,9,43). Specifically, the extent of the change in the force of infection (*θ*_*λ*_) represents the strength of the interaction effect, such that values above 1 indicate positive interactions (e.g., *θ*_*λ*_ = 2 implies a doubling of the force of infection), values below 1 indicate negative interactions (e.g., *θ*_*λ*_ = 0.5 represents a halving of the force of infection, while *θ*_*λ*_ **=** 0 represents complete inhibition), and a value of 1 indicates no interaction. Meanwhile, the time spent in compartment T (*7*/*δ*, where *δ* is the rate at which individuals leave the T compartment) represents its duration. A model schematic can be seen in Figure 1, and the full model equations can be found in the Supplementary Materials (S1 Text). We have previously used a similar model to infer interactions between influenza and RSV (3).

**Figure 1.**
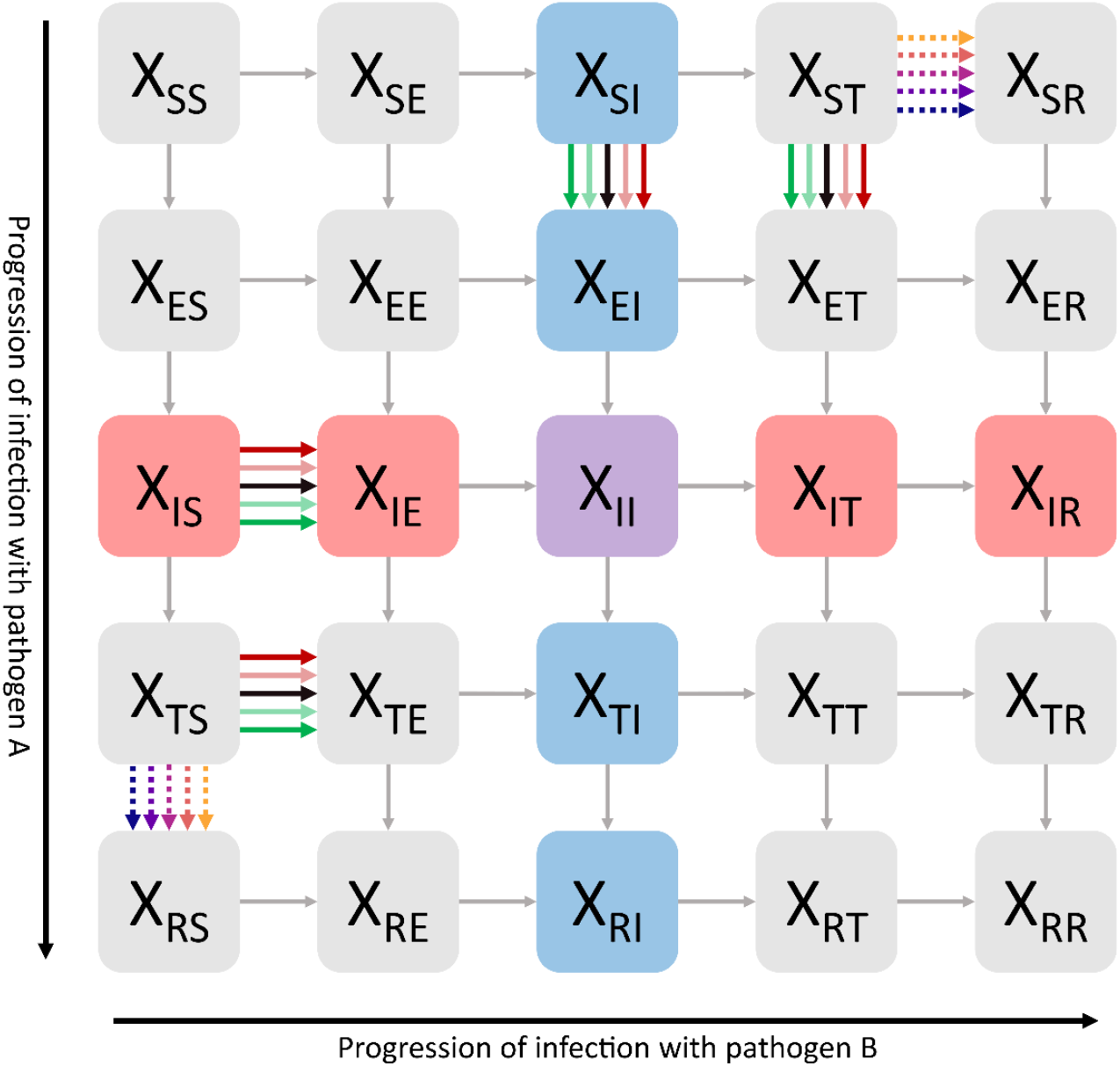
Schematic of the model used to generate synthetic data. Each box represents a distinct model state, with the first subscript indicating infection status with respect to pathogen A, and the second subscript indicating status with respect to pathogen B. Infection with pathogen A is indicated in red and progresses vertically, while infection with pathogen B is indicated in blue and progresses horizontally; coinfection is shown in purple. Arrows represent possible transitions between states. Transitions impacted by changes in interaction strength are shown in a gradient of colors from green to red, while transitions impacted by changes in interaction duration are shown as dotted arrows with a gradient of colors as in Figures 2, 4, and 5.

We parameterized the model such that pathogen A was based on influenza and pathogen B was based on RSV, two viruses between which a moderate-to-strong, negative interaction is likely to exist (3,4). Where information was available, parameter values for each of the two pathogens were based on past observational and modeling studies of both pathogens. Otherwise, parameter values were chosen such that, in the absence of an interaction effect, simulations reproduced patterns resembling real-world influenza and RSV outbreaks in temperate climates (i.e., annual periodicity with a single peak in winter). Specific parameter values used, as well as sources where available, can be found in Table 1.

**Table 1.**
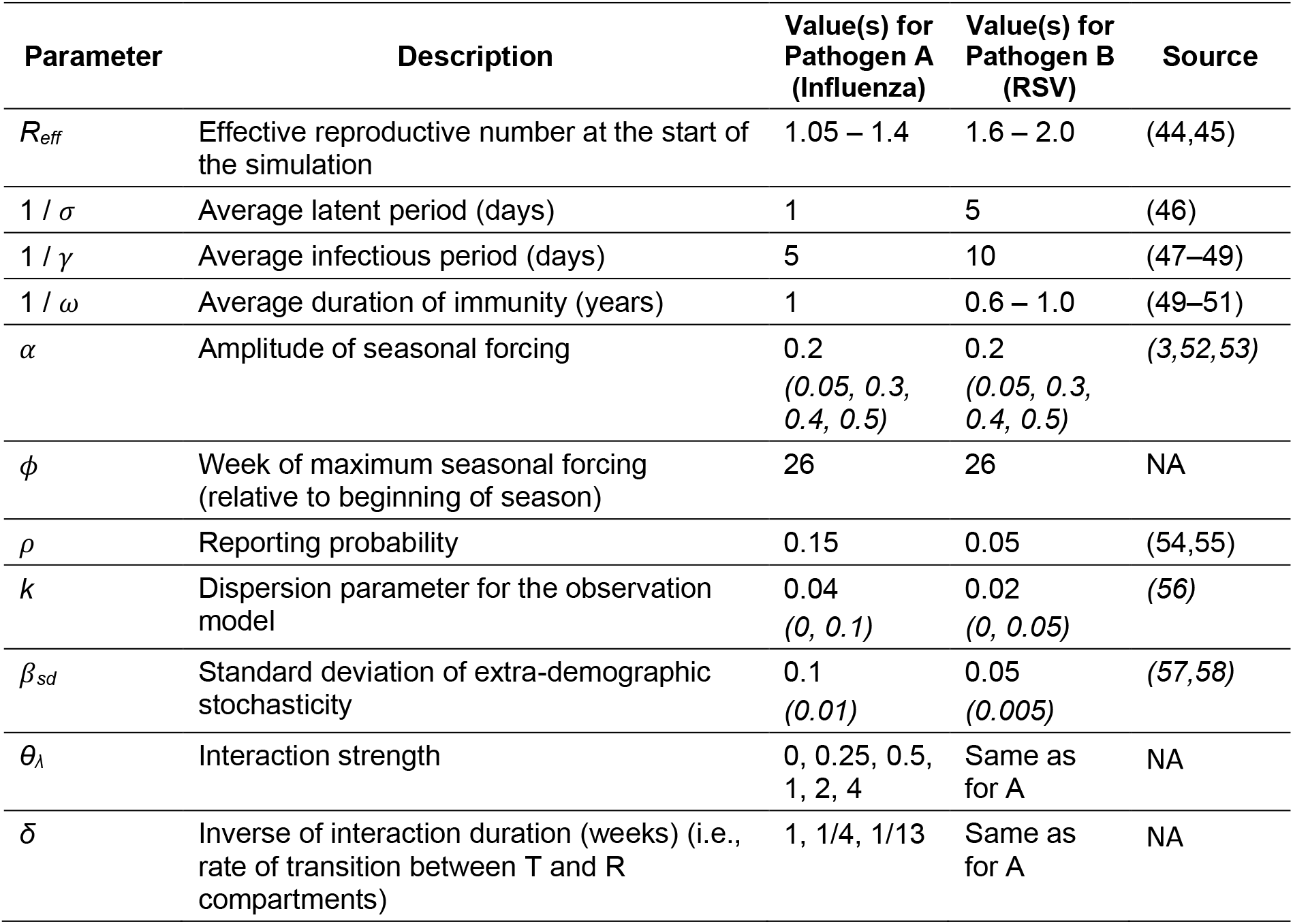
Parameters and parameter values used to generate synthetic data. Ranges indicate parameters that were allowed to vary in the main analysis. Values used for sensitivity analyses are shown in parentheses and italics.

Influenza and RSV display similar seasonal patterns, with both viruses circulating primarily during the winter in temperate regions (59). Indeed, some work suggests that influenza viruses and RSV respond similarly to several environmental drivers, including temperature, humidity, and rainfall (18,60–65). To capture this, we introduced a common seasonal driver to the transmission rate of both pathogens, such that:

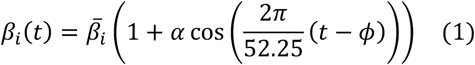

where *β*_*i*_ (*t*) is the transmissibility of pathogen *i* at time *t*, 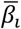 is the yearly average transmissibility of pathogen *i, α* is the seasonal amplitude of transmissibility, and *ϕ* is the week during which transmissibility is highest. The division by 52.25 inside the cosine function imposes a yearly periodicity on the transmission rate, and accounts for the weekly timescale of our synthetic data. This shared seasonal forcing represents a potential confounder when estimating the interaction effect between the two pathogens.

Unlike RSV, which is comparatively antigenically stable, influenza viruses undergo rapid “antigenic drift” (66). Furthermore, there is year-to-year variation in which influenza subtype(s) drive seasonal outbreaks. To capture the impact of this antigenic diversity on population-level susceptibility, we incorporated yearly “surges” in immunity loss into the model of influenza only, as in (67). Specifically, these surges occurred on average during the thirteenth week of each season, and could vary in size from 5-30% of the recovered population (see S1 Text for additional details). The model population was set to 5 million, and the birth and death rates were both set to 2×10^-4^ per week (i.e., 10.45 births per 1000 people per year), such that the population size remained constant over time.

#### Observation Model

The SEIRSxSEIRS model describes the true number of individuals in each model state at each timepoint. However, real-world surveillance systems cannot perfectly capture the true number of cases of a disease at any given time. Influenza and RSV in particular often cause mild or asymptomatic illness, such that many infected individuals do not seek healthcare and therefore are not included in surveillance data. For this reason, all model states in the transmission model (Figure 1) are latent, or unobserved. In order to produce synthetic data that are representative of real-world surveillance data, an observation model is needed to describe how observed data arise from the underlying latent process.

Here, we generate synthetic data by drawing observed cases from a negative binomial distribution each week. Specifically, the distribution at time *t* has mean *μ* = *ρ*_*i*_*H*_*i*_(*t*), where *ρ*_*i*_ is the probability that a case of pathogen i is reported, and *H*_*i*_ is the number of cases of pathogen i who have recovered between timepoints *t-1* and *t* (i.e., we assume that cases report near the end of the infectious period), such that *dH*_*A*_ /*dt* is equal to (*X*_*IS*_ + *X*_*IE*_ + *X*_*II*_ + *X*_*IT*_ + *X*_*IR*_)*γ*_*A*_ and *dH*_*B*_/*dt* is equal to (*X*_*SI*_ + *X*_*EI*_ + *X*_*II*_ + *X*_*TI*_ + *X*_*RI*_)*γ*_*B*_ (see also Equation 1 in S1 Text). The observation model has dispersion parameter *k* (i.e., the distribution has variance *σ*^2^ = *μ* + *kμ*^2^), such that higher *k* yields higher variability in reporting.

### Generation of Synthetic Data

Synthetic data were generated using a wide range of interaction strengths (*θ*_*λ*_ = 0, 0.25, 0.5, 1, 2, 4) and durations (*7*/*δ* = 1, 4, or 13 weeks), including both positive and negative interactions, as well as simulations with no interaction between the two pathogens. All interactions were symmetric with regard to both strength and duration; in other words, the impact of pathogen A on pathogen B was of the same strength and duration as the impact of B on A.

Here, we focus specifically on short-term interactions, as these are more realistic for unrelated pathogens where adaptive cross-immunity is unlikely, and our model is parameterized to produce outbreaks similar to those of influenza and RSV. While long-term interactions due to adaptive immune mechanisms can occur between related pathogens (10,68,69), understanding these interactions is a somewhat distinct problem, where knowledge of the underlying mechanisms (e.g., cross-immunity between paramyxoviruses (10), antibody-dependent enhancement between dengue serotypes (69)) may provide some prior information about the interaction’s sign and duration. However, a test of CCM for long-term interactions can be found in (36).

For each strength-duration pair, we generated 100 synthetic datasets. The timing and magnitude of surges in immune loss (described above) were allowed to vary in each of these 100 simulations, as were values of *R*_*effA*_, *R*_*effB*_, *ω*_*B*_ (see S1 Text for details). This allowed for the generation of some datasets where pathogen A tended to peak before pathogen B, and others where pathogen B tended to peak before pathogen A, a factor we have found in past work to be important in determining whether an interaction can be accurately characterized based on data. The same 100 sets of values for *R*_*effA*_, *R*_*effB*_, *ω*_*B*_, and the timing and magnitude of the surges in immune loss were used when generating datasets for each of the eighteen combinations of interaction strength and duration.

All datasets were generated using a stochastic model implementation. We incorporated both demographic and extra-demographic (or environmental) stochasticity, with extra-demographic stochasticity modeled by multiplying the transmission rate of each pathogen by gamma white noise with standard deviation equal to *β*_*sd*_ at each timestep (see Table 1) (70,71). The observation model (see above) contributes additional stochasticity.

All simulations were first run for 20 years to allow the system to approach equilibrium. The model was then run for a further 10 years; these final 10 years of data were used in the analyses described below. To prevent the extinction of pathogen A, we added 100 individuals infected with pathogen A to the system from outside the model population each year during week 14. A representative dataset, where we have varied only the strength and duration of the interaction, can be seen in Figure 2A; further examples can be found in Figure S1.

**Figure 2.**
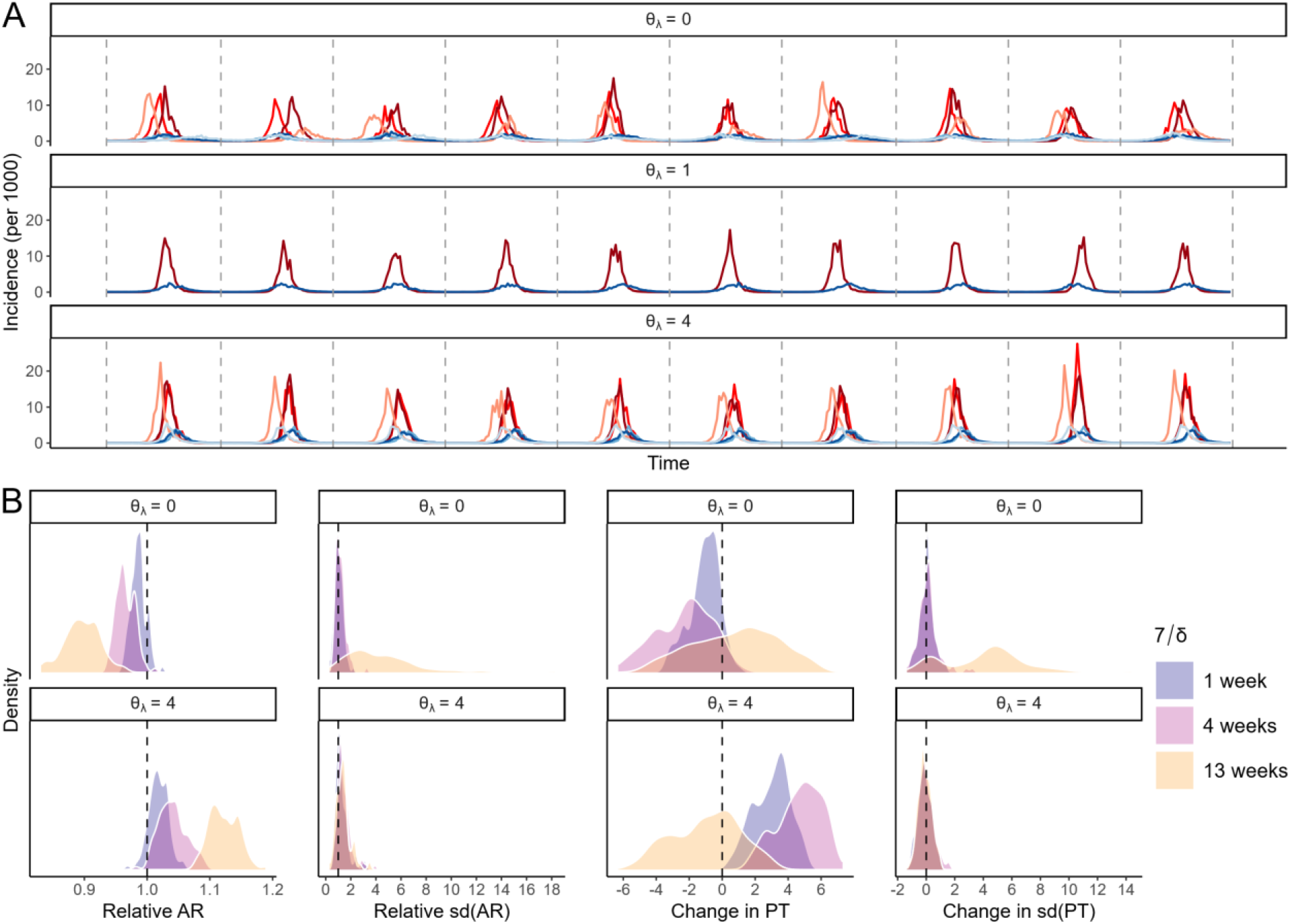
Impact of interactions on outbreak dynamics. (A) Example data consisting of ten years of simulated observations for pathogen A (red) and pathogen B (blue), generated with varying interaction strength (*θ*_*λ*_) and duration (7/*δ*) and all other parameters held constant at the values shown in Table 1 (here, *Ri*_*1*_ = 1.38, *Ri*_*2*_ = 1.89, 1/*ω*_*2*_ = 49.5 weeks). Specifically, data are shown for simulations with a strong negative interaction (top), no interaction (middle), and a strong positive interaction (bottom); line colors show the duration of the interaction, with the darkest colors indicating a duration of 1 week, medium colors indicating duration 4 weeks, and the lightest colors indicating duration 13 weeks. Dotted vertical lines indicate the beginning of each new epidemic season, with each season lasting one year. (B) The distribution of the change in attack rate (i.e., the total number of observed cases in a single year), year-to-year variation (i.e., standard deviation across all years) in attack rate, peak timing (i.e., the week during which observed incidence for a given year is maximal), and year-to-year variation (i.e., standard deviation across all years) in peak timing of pathogen B when either a strong negative (top) or strong positive (bottom) interaction is at play, relative to the dynamics when no interaction is present. Shading indicates the duration of the interaction; dotted vertical lines indicate no change.

In addition to the main analysis, we conducted several sensitivity analyses in which we varied 1) the number of years of data used for inference (20 vs. 10), 2) the amplitude of seasonal forcing (*α* = 0.05, 0.3, 0.4, 0.5), and 3) the amount of process 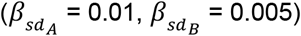 and observational (*k*_*A*_ = 0, 0.1; *k*_*B*_ = 0, 0.05) noise.

The transmission model was coded and run using the package pomp (version 6.1) (72).

### Real-World Data

In addition to synthetic data, we applied all methods to observed influenza and RSV positivity data from Hong Kong (73) and Canada (74). These data were chosen because we have previously fit a mechanistic transmission model to data from these two locations, and found evidence of a negative interaction in both datasets (3), allowing us to assess whether each method tested here yields similar results.

The data are more comprehensively described and visualized in (3). Briefly, data consist of the weekly number of tests positive for influenza and for RSV, as well as the total weekly number of tests conducted, over the course of four to six seasons. Samples were primarily obtained from inpatients and emergency department patients. From this data we calculated the weekly proportion of tests positive for influenza and for RSV, which we used as our case data when applying the statistical methods detailed in the following section. Note that, for Canada, we combined all (sub)types of influenza (H1N1, H3N2, and B); for Hong Kong, however, where H3N2 outbreaks often lead to multiple yearly peaks, we included only H1N1 and B, as in (3). For methods that allowed it, we controlled for weekly mean absolute humidity, calculated from the US National Centers for Environmental Information’s Global Surface Summary of the Day (GSOD) data (75), which we obtained using the R package “GSODR” (76), using the Clausius-Clapeyron relation (77). Since the Canadian surveillance data cover the entire country, we used the median absolute humidity across all available stations each week. Absolute humidity data were chosen due to the likely influence of humidity on the transmission of both influenza and RSV (18,61,64).

### Statistical Methods

We tested five non-mechanistic methods: (1) Pearson correlations, (2) generalized additive models (GAMs), (3) Granger causality, (4) transfer entropy, and (5) convergent cross-mapping (CCM). Characteristics of these methods are summarized in Table 2. Briefly, we applied each method to all 100 datasets for each of the 16 interaction parameter combinations listed above (see “Generation of Synthetic Data”), as well as to the surveillance data from Hong Kong and Canada. To fulfill the assumptions of the first three methods in particular, we log-transformed and centered the data prior to analysis (30); for consistency, this was done for all five methods tested. For methods accounting for shared seasonality, the magnitude of seasonal forcing at each timepoint was similarly transformed. Finally, we checked whether each synthetic dataset was stationary using both the Augmented Dickey– Fuller (78) and Kwiatkowski-Phillips-Schmidt-Shin (KPSS) (79) tests. Nonstationary datasets (n=2 of 1800) were removed from consideration.

**Table 2.**
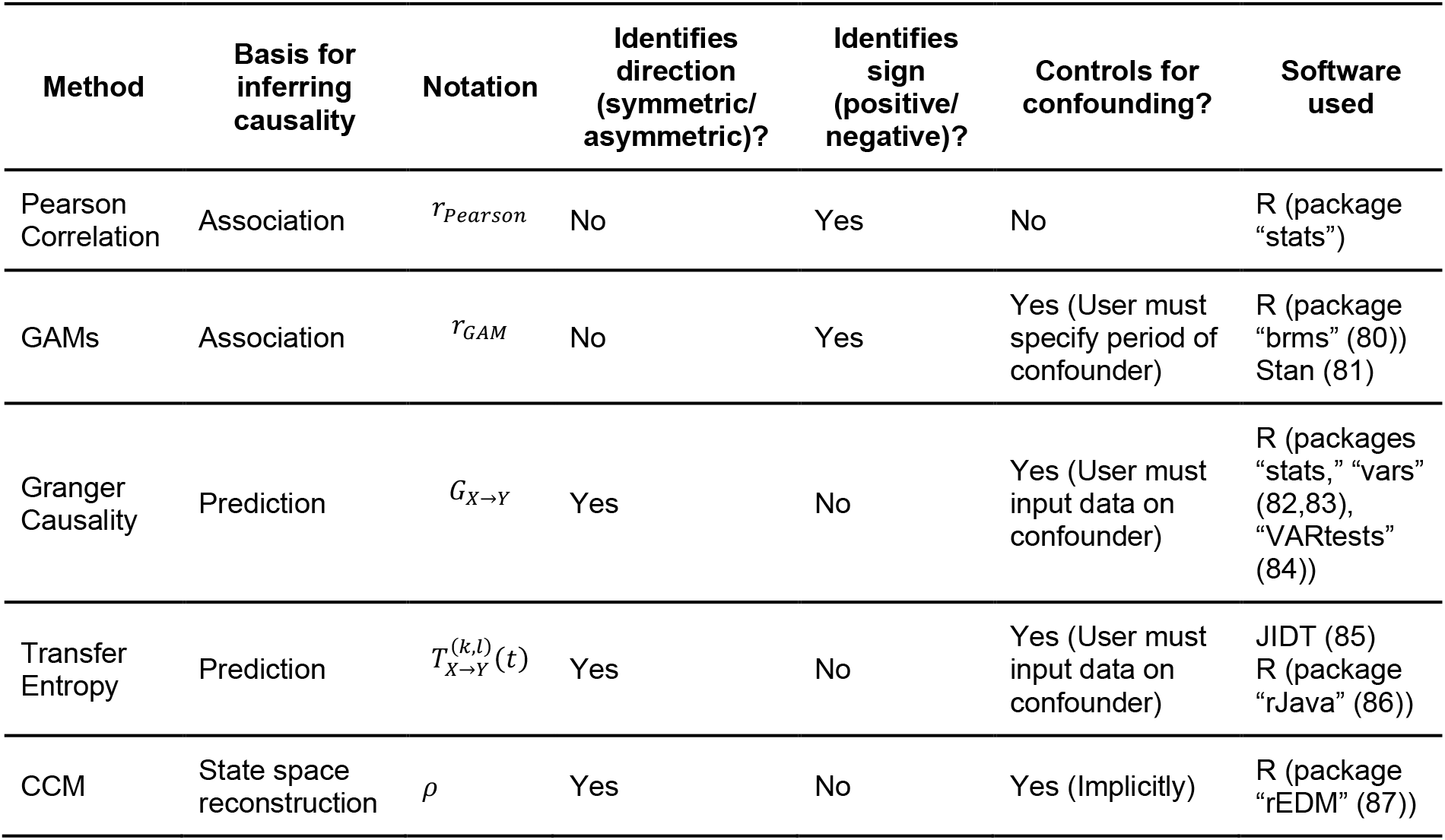
Overview of all non-mechanistic methods evaluated.

A method was said to have identified an interaction for a given dataset if the results were statistically significant (p < 0.05). Meanwhile, the effect estimates were evaluated as indicators of the relative strength of the interactions; for Pearson correlations and GAMs, effect estimates also indicated sign (i.e., estimates could be positive or negative). We describe this process in detail for each method in the following subsections.

#### Pearson Correlation

The Pearson correlation coefficient measures the strength of the linear association between two datasets, and is defined as:

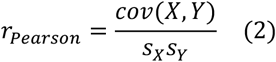

where *cov*(*X, Y*) is the covariance of *X* and *Y, s*_*X*_ is the standard deviation of *X*, and *s*_*Y*_ is the standard deviation of *Y*. Values above 0 indicate a positive association, while values below 0 indicate a negative association; values of exactly 1 or –1 indicate a perfect linear relationship between the two datasets. Note that this method cannot be used to test directionality, i.e., *r*_*XY*_ = *r*_*YX*_.

We calculated the Pearson correlation coefficient between the time series of our two modeled pathogens for all synthetic datasets. Point estimates and two-tailed p-values were obtained using the cor.test function in the package “stats” (version 4.4.0). Unlike many of the other methods tested here, correlation coefficients can be negative, allowing us to evaluate the extent to which interaction sign is correctly inferred. This approach is simple and does not account for potential confounding due to the shared seasonal driver. Furthermore, as we have emphasized in the introduction above, the complexity and nonlinearity inherent to infectious disease dynamics can lead to unintuitive relationships between observed case counts. However, although we do not necessarily expect this method to perform well, we have included it because correlation- and regression-based methods continue to be used to characterize pathogen-pathogen interactions (as detailed in the Introduction), and, to our knowledge, the accuracy of this approach has yet to be formally evaluated.

#### Generalized Additive Models

Unlike in standard regression, where an outcome of interest is fit using the raw values of a set of predictors, generalized additive models (GAMs) fit the outcome data to smooth functions of the predictors (25,88). By using a penalized likelihood approach, GAMs ensure that these smooths contain enough “wiggliness” to fit the data, but not so much as to cause overfitting. This allows GAMs to flexibly describe nonlinear relationships between variables. It is this ability that makes GAMs a potentially attractive approach for identifying interactions: by fitting the seasonal cycles in the data, we can explore the association between the time series after removing the effects of shared seasonal forcing. Because our synthetic data represent case counts and not the underlying force of infection, this method does not perfectly control for confounding; nonetheless, it offers a potential advantage over the correlation coefficient method discussed above.

For each synthetic dataset, we fit a bivariate normal GAM to the time series of case counts for both pathogens, with a smooth on week of the year (i.e., 1 through 53; WOY) as the only predictor. When fitting GAMs, it is necessary to specify the basis dimension for any smooths, which sets an upper bound on the flexibility of the curve (25,88). For WOY, we set this value to 53. Furthermore, we used a cyclic cubic regression spline, which ensures that the smooth is continuous at the beginning and end of the year (25). Thus, the fitted model for each dataset was of the form:

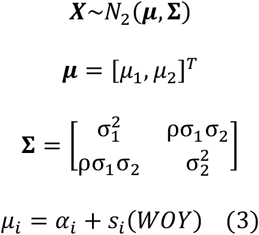

where ***X*** *=* [*X*_*1*_, *X*_*2*_]^*T*^ is a matrix containing case counts for the full study period for both pathogens, which is modeled as a bivariate normal distribution with mean ***μ*** and covariance matrix ***Σ***. The mean and standard deviation for each pathogen *i* are indicated by *μ*_*i*_ and *σ*_*i*_, respectively, while *ρ* is the correlation coefficient between the two time series. In the final equation, “s” indicates a smooth term of the specified predictor; *α*_*i*_ represents the pathogen-specific intercept.

All GAMs were fit using a Bayesian approach as implemented by the “brms” package (version 2.22.0) (80), which interfaces with the probabilistic programming language Stan (81). For each pair of time series, we ran 2000 warmup iterations, followed by 1000 sampling iterations. In addition to the relationship between the predictor and outcome variables, the bivariate normal model also fits the residual correlation between predictors. We took the median of the posterior distribution of these correlation coefficients to be our point estimate of interaction strength. We constructed 95% confidence intervals by calculating the 2.5th and 97.5th percentiles of this distribution, and estimates were considered to be significant if this interval did not contain 0. As with correlation coefficients calculated from the raw data, this method is incapable of distinguishing between symmetric vs. asymmetric interactions. Datasets where fitting these models resulted in 1) any divergent transitions after warmup, or 2) lack of convergence, as indicated by an R-hat of greater than 1.05 for any model parameter, were removed from consideration before further analysis. Additional methodological details can be found in S1 Text.

#### Granger Causality

The previous two methods rely on correlations between the time series of two pathogens to determine whether an interaction is present, which, as discussed above, is potentially problematic. Wiener-Granger causality (Granger causality for short) (21,22) instead infers causation based on predictive ability. Specifically, Granger causality measures the extent to which knowledge of the past values of *X* improves the prediction of *Y* above and beyond the predictive capability of past values of *Y* (and any confounders of interest) alone. Here, we calculate this quantity using vector autoregressive models (VARs), as in Granger’s original implementation (21), although this framework can be extended to more complicated time series models (89,90). To account for shared seasonality, we calculated the true extent of seasonal forcing over time according to Equation 1, and included this time series as a predictor in all models. We therefore test for Granger causality by comparing the following two models:

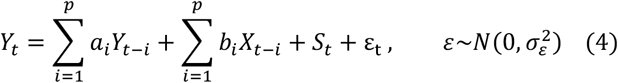

and

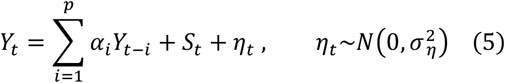

where *p* is the order of the autoregressive models, and *S* is the seasonal forcing component of Equation 1 (i.e., everything except 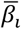, which is pathogen-specific). The terms *ε*_*t*_ and *η*_*t*_ represent the remaining error in *Y*_*t*_ not explained by the models, which are assumed normally distributed with variance equal to 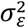 and 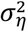, respectively. If 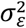 is significantly less than 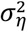, it is said that *X* “Granger causes” *Y*. Despite this terminology, we emphasize that forecasting ability does not necessarily indicate that one variable has a causal effect on another. Thus, *X* may “Granger cause” *Y*, but may not actually have a causal effect on *Y* (91,92).

To apply this method to our synthetic datasets, we first identified the ideal order for each synthetic dataset as the value that minimized the BIC of the fitted VAR models. We allowed for a maximum order of 20, the sum of the upper bounds of the generation time for both pathogens (2 weeks for pathogen A and 5 weeks for pathogen B) and the maximum interaction duration tested (13 weeks) (see S1 Text for a derivation of these values). We also conducted a sensitivity analysis using a larger value (see S1 Text). The magnitude of the estimated effect of one pathogen on the other was then calculated as initially suggested by (93):

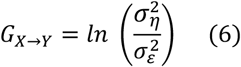

These values were deemed to be statistically significant if any elements of *b*_*i*_ were non-zero, as evaluated by an F-test.

All analyses were conducted using the “stats” (version 4.4.0), “vars” (version 1.6.1) (82,83), and “VARtests” (version 2.0.5) (84) packages; a link to the full code used can be found under “Implementation” above. Note that, unlike the above methods based on correlations, Granger causality allows for the effect of pathogen A on pathogen B to differ from the effect of pathogen B on pathogen A, meaning that asymmetric interactions can theoretically be identified. However, the effect estimate calculated in Equation 6 is always positive; thus, Granger causality is incapable of distinguishing between positive and negative interactions.

#### Transfer Entropy

Like Granger causality, transfer entropy (23,94) estimates the extent to which past information on *X* informs the current state of *Y*, above and beyond past information on *Y* itself. However, rather than VARs and other parametric time series models, transfer entropy instead relies on ideas from information theory (95). Specifically, the transfer entropy from *X* to *Y*, conditional on the shared seasonal forcing, is defined as:

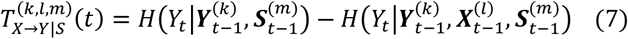

Here, *k, l*, and *m* are the history lengths (i.e., the number of timepoints included in the analysis) for the target (*Y*), source (*X*), and seasonal forcing (*S*) data, respectively. Meanwhile, 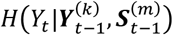 is the conditional Shannon entropy (96) of *Y*_*t*_ given 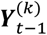 and 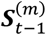, a measure of the amount of uncertainty in the value of *Y* at time *t*, after accounting for the previous history of *Y* up to *k* timepoints in the past and *S* up to *m* timepoints in the past. The term 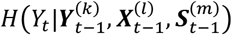 is similar, this time also conditioning on the past history of *X* up to timepoint *t – l*. More specifically, Shannon entropy measures uncertainty by accounting for both the range of possible values taken by *Y*_*t*_, as well as the probability of each value, such that variables with a narrow possible range of values will have low entropy (i.e., low uncertainty), whereas variables with an equal probability of taking a wide range of values will have high entropy. Notably, Equation 7 is equivalent to the conditional mutual information between *Y*_*t*_ and 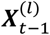 given 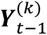 and 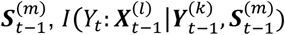, a measure of the reduction in uncertainty about the value of *Y* at time *t* due to knowledge of the past history of *X*, after accounting for the past history of *Y* and *S* (95). Like with Granger causality, values above zero (i.e., where the uncertainty in *Y* at time *t* when accounting for the past histories of *X, Y, S* is lower than the uncertainty in *Y* at time *t* when accounting only for the past histories of *Y* and *S* alone) are taken to imply that *X* causes *Y*.

Overall, transfer entropy may be viewed as a nonparametric alternative to Granger causality (94,95,97), and for Gaussian variables, the two are equivalent (98). Because VARs may not perform well when dynamics are highly nonlinear (99), and data transformations can greatly affect the resulting Granger causality estimate (100), transfer entropy may be a more appropriate approach in these cases. However, calculating transfer entropy is more computationally intensive than running the relatively simple VARs necessary to calculate Granger causality (95).

We calculated the transfer entropy for all datasets using the Kraskov-Stögbauer-Grassberger (KSG) technique with four nearest neighbors (101), as implemented by the Java Information Dynamics Toolkit (JIDT) (85). The history length for the target (i.e., outcome) data was fixed to 2 for pathogen A and 5 for pathogen B, based on the generation times for influenza and RSV, respectively; the history length for the source (i.e., predictor) data was set to 20 for both pathogens (see S1 Text). As with Granger causality, we also performed a sensitivity analysis where larger values were used (S1 Text). For the shared seasonal forcing, history length was set to 1. Due to the range of interaction durations modeled, we also allowed for lags greater than 1 week between *X* and *Y*; specifically, we tested lags of 1, 2, 4, and 13 weeks. Only results for the lag yielding the highest estimates of transfer entropy are shown (102). Significance was assessed by generating 500 permutations of the data under the null hypothesis, and comparing our point estimates to estimates calculated from these surrogate datasets (85).

Analyses were conducted using JIDT (version 1.6.1) (85) and the R package “rJava” (version 1.0.11) (86). Like Granger causality, point estimates for transfer entropy are always positive, and therefore do not allow for the characterization of positive vs. negative interactions.

#### Convergent Cross-Mapping

The above methods make sense for systems where *X* and *Y* contain unique information, and therefore the predictability of *Y* in the absence of information about *X* can be assessed simply by removing the time series of *X* from consideration. However, in nonlinear dynamical systems driven by an underlying deterministic skeleton, a time series *Y* will itself contain information on the past values of its cause *X*. In such situations, the utility of Granger causality and transfer entropy is controversial (24,30).

Convergent cross-mapping (CCM) is a relatively new approach that takes advantage of this inherent lack of independence between time series (24). Specifically, to determine whether *X* causes *Y*, CCM uses *L* lagged values of *Y* to construct a shadow manifold, *M*_*Y*_; this is done according to Takens’ theorem (103), which states that information about all states/the state space of a dynamical system can be reconstructed based on the lagged values of a single state of that system. Then, the method assesses whether the values of *X* can be estimated based on the topology of *M*_*Y*_. In other words, unlike Granger causality and transfer entropy, which infer that *X* causes *Y* if past values of *X* improve predictions of some current value of *Y*, CCM infers that *X* causes *Y* if current values of *Y* can be used to “predict” current or past values of *X*. Crucially, for causation to be demonstrated, the quality of these predictions must increase with increasing *L*, or the amount of data available for building the shadow manifold (i.e., there must be convergence).

To perform CCM, we first chose the ideal embedding dimensions (i.e., the number of lags used to construct the shadow manifolds) for each synthetic dataset by determining which values yielded the most accurate within-pathogen predictions for each pathogen. We allowed a maximum embedding dimension of 2 for pathogen A and 5 for pathogen B (see S1 Text). We also allowed for a lagged effect of one pathogen on the other by selecting the negative lag that yielded the highest cross-map skill (104); a maximum lag of 20 weeks was permitted (see S1 Text). As for the previous two methods, we also ran a sensitivity analysis using larger values for the maximum embedding dimensions and lags (S1 Text). The selected values for the embedding dimensions and lags were then used to run CCM with 100 samples for each value of *L*. Point estimates were taken to be the median cross-map skill across all samples for the largest *L*. We note that, since CCM is expected to yield null results for two variables that are not causally linked but have a common environmental driver (24), there is no need to explicitly account for the shared seasonal forcing between pathogens when applying this method.

We tested two methods of assessing significance. The first approach considers a result to be significant so long as there is convergence, as in CCM’s original implementation (24). Specifically, we use a nonparametric bootstrap approach to determine whether the cross-map skill for the maximum value of *L* is significantly greater than the cross-map skill for the minimum value of *L*, as in (36). The second approach, originally suggested by (18), compares the cross-map skill obtained from the data for the maximum value of *L* to a null distribution of cross-map skill calculated using seasonal surrogates of the hypothesized causal pathogen’s data. Specifically, we generate 500 surrogate time series by maintaining the seasonal pattern in the data, but reshuffling the residuals. A p-value is then calculated as:

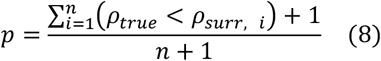

where *ρ*_*true*_ is the cross-map skill calculated from the data, *ρ*_*surr,i*_ is the cross-map skill calculated using surrogate time series *i*, and *n* is the number of seasonal surrogates considered (30,105).

Compared to the first method, this approach may be better suited to systems where shared seasonality presents a challenge to causal inference (18).

Analyses were conducted using the R package “rEDM” (version 1.15.4) (87). As with Granger causality and transfer entropy, cross-map skill is always greater than zero. For this reason, CCM does not allow for positive and negative interactions to be differentiated (although similar methods exist that use scenario exploration to characterize the sign of the effect of one variable on another (18,106)).

### Performance Assessment

We assessed the overall performance of each method by calculating the sensitivity (i.e., the proportion of datasets where the presence of an interaction is correctly inferred) and specificity (i.e., the proportion of datasets where the absence of an interaction is correctly inferred). Here, we considered results using Pearson correlation coefficients and GAMs to be accurate only if they correctly identified the sign of the interaction; for all other methods, which only yield positive point estimates, accuracy was assessed on the basis of significance alone. While this decision technically disadvantages correlation coefficients and GAMs, we feel that it is also important to assess the accuracy of the extra information these methods capture about interaction sign.

To determine whether the true underlying interaction parameters influenced accuracy, we also calculated the percentage of synthetic datasets correctly classified by each method for each combination of interaction parameters. Again, for Pearson correlation coefficients and GAMs, we considered both significance and sign in assessing accuracy.

If the tested methods are correctly identifying the relative strength of the modeled interactions, we expect that the methods will yield larger point estimates for datasets where the strength of the underlying interaction is greater. To assess this, we fit a linear mixed-effects regression model on the point estimates obtained from each method, using true strength, true duration, and the interaction between them as predictors, as well as a random intercept on the parameter set. To account for the fact that point estimates from some methods are necessarily positive, we took the reciprocal of true interaction strength for negative interactions prior to fitting; interactions with strength 0 (i.e., complete inhibition), for which the reciprocal is undefined, were assigned a value of 16; results remained qualitatively similar when different values were chosen. We also log2-transformed both the absolute value of the point estimates, and the true interaction strengths, such that the regression models captured the relative change in outcome resulting from a doubling of interaction strength. This was done because the range of point estimates produced varied by method.

## Results

### Symmetric interactions between pathogens led to observable differences in epidemic dynamics

Changing the strength and duration of the interaction between our two modeled pathogens led to observable differences in the dynamics of both pathogens, particularly for long-lasting interactions (Figure 2A, Figure S1). This suggests that outbreak data, which are collected at the population level, may nonetheless contain signal of individual-level interactions. When holding all other model parameters constant, negative interactions generally reduced both the average attack rate and the average peak week of pathogen B. In contrast, positive interactions had the opposite effect (Figure 2B). However, this was not always the case: long-lasting negative interactions could push outbreaks later, and vice versa. Furthermore, while the timing and size of outbreaks of pathogen B were typically highly consistent across seasons, long-lasting negative interactions greatly increased year-to-year variability. Similar patterns were observed for pathogen A (Figure S2). These findings illustrate a key point from the introduction: although interactions can impact infectious disease transmission, these effects can be subtle and their direction unintuitive.

### Non-mechanistic methods applied to incidence data frequently fail in determining whether or not an interaction between two pathogens exists

The overall accuracy of all methods at identifying the presence or absence of an interaction effect is displayed in Figure 3. For Pearson correlation coefficients and GAMs, negative interactions were only considered to be correctly inferred if the resulting point estimate was also negative; for all other methods, which can only yield positive point estimates, results were considered accurate for negative interactions as long as point estimates were significantly greater than zero. For Granger causality and transfer entropy, only the results of tests accounting for seasonality are shown (see Figure S3 for results ignoring shared seasonality). Of the tested methods, GAMs performed best, with 85.2% sensitivity and 72.5% specificity. All other methods showed much worse performance: while sensitivity was often high, specificity was below 50% for all other methods. In other words, among simulations where no interaction was modeled, all methods other than GAMs yielded false positives more often than not. Results were no better when larger values were chosen for various method parameters (e.g., lags, history lengths, and embedding dimensions), and for CCM in particular, using larger embedding dimensions and lags often led to even lower specificity (Figure S4). For most methods capable of distinguishing interaction direction (Granger causality, transfer entropy, and CCM using seasonal surrogates), false positives were more common when trying to classify the interaction effect of pathogen B on pathogen A. CCM using convergence as the criterion for significance was the only exception, but improved specificity came at the cost of sensitivity. Overall performance quality was similar for these three methods; as expected (20,107), Pearson correlation coefficients were by far the least accurate.

**Figure 3.**
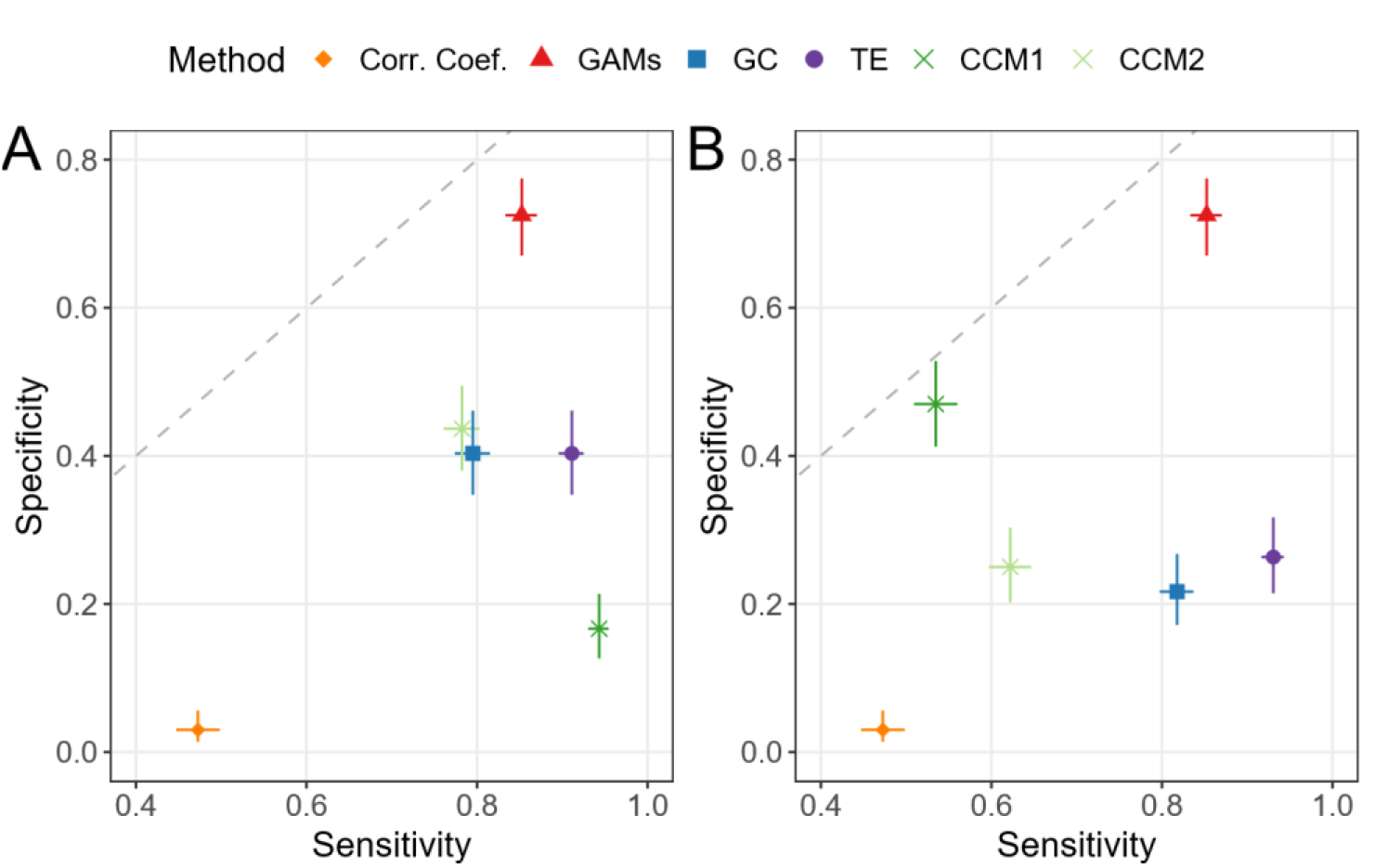
Sensitivity and specificity of all tested methods. Results for the interaction effect of pathogen A on pathogen B are shown in (A), while results for the effect of pathogen B on A are shown in (B). Method is indicated by both point shape and color; CCM1 refers to the method assessing significance based on convergence, while CCM2 refers to the method assessing significance using seasonal surrogates. The crosshairs on each point indicate 95% confidence intervals, obtained using a binomial test. The dashed diagonal line shows where sensitivity is equal to specificity. Note that the x-axis begins at 0.4. For correlation coefficients and GAMs, negative interactions were only considered to be correctly detected if the associated point estimate was also negative; all other methods cannot distinguish between positive and negative interactions, and so were considered to have correctly identified an interaction if the point estimate was significantly above For Granger causality and transfer entropy, only results for implementations controlling for shared seasonality are shown.

### Accuracy was dependent on the true strength and duration of the interaction

Accuracy for all methods by true strength and duration of the modeled interaction is plotted in Figure 4. In general, accuracy was higher for interactions with longer duration, perhaps due to increased signal of longer interactions in the data, especially when interaction sign was negative. Accuracy was typically similar for positive and negative interactions, although CCM performed better for positive interactions, at least when the effect of pathogen B on pathogen A was being tested (Figure 4H, J), and correlation coefficients performed particularly poorly for negative interactions (Figure 4A). Interaction strength had little effect on accuracy for all of the tested methods.

**Figure 4.**
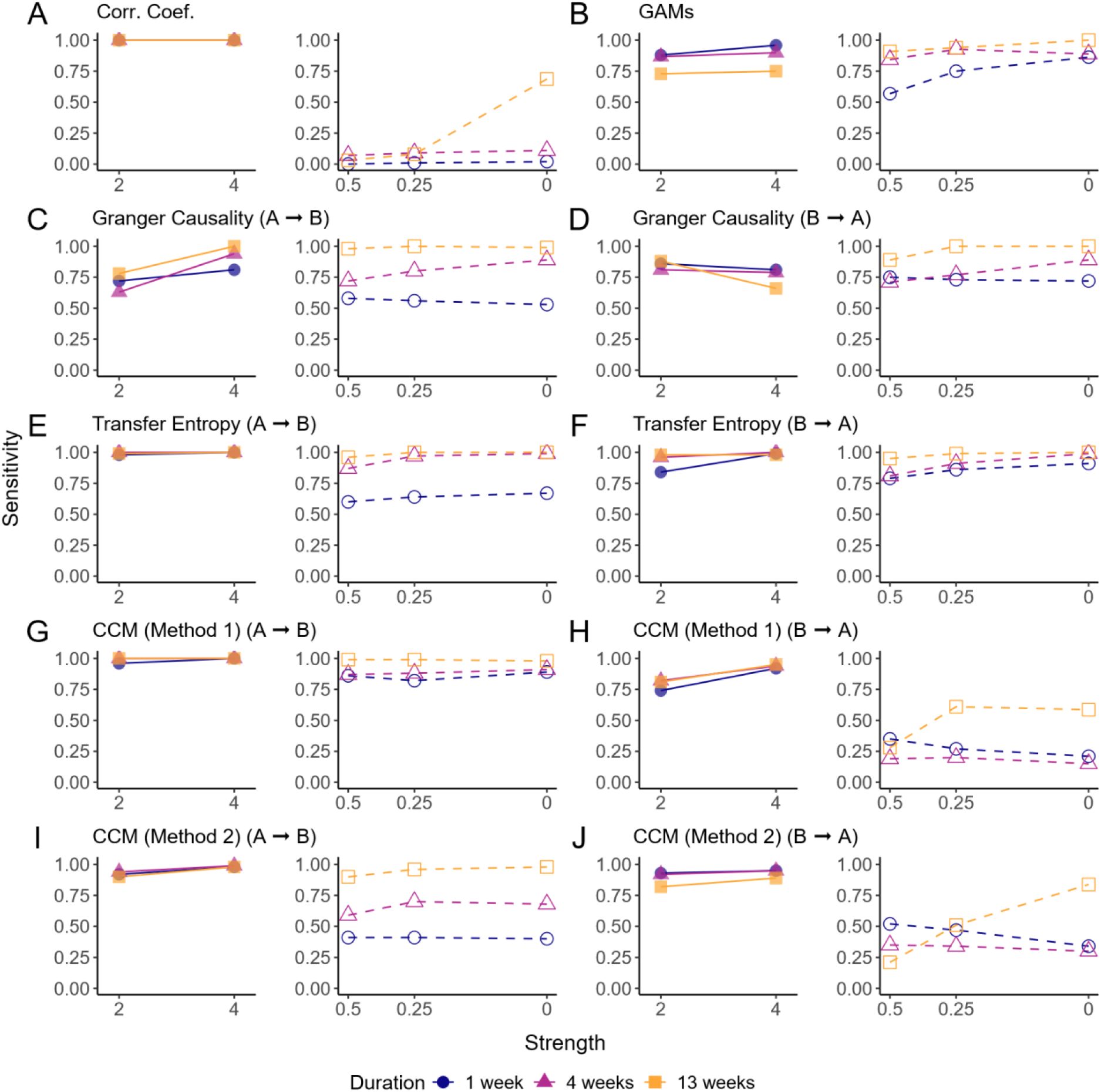
Sensitivity of all methods by true interaction strength and duration. Results are shown for correlation coefficients (A), GAMs (B), Granger causality (C and D), transfer entropy (E and F), CCM using convergence to assess significance (G and H), and CCM using seasonal surrogates to assess significance (I and J); for methods capable of distinguishing directionality, results for the effect of pathogen A on pathogen B are shown in (C), (E), (G), and (I), while results for the effect of B on A are shown in (D), (F), (H), and (J). For each method, results for positive interactions are shown in the left panel, and results for negative interactions are shown on the right. True interaction strength is shown on the x-axis, arranged from weaker interactions on the left to stronger interactions on the right. Results from simulations generated with positive interactions are displayed as filled points and connected with solid lines, while results from simulations with negative interactions are displayed as hollow points and connected with dotted lines. Point and line color indicate the true interaction duration.

### Magnitude of the point estimates was not consistently associated with true interaction strength

In addition to identifying whether an interaction between two pathogens exists, methods should ideally provide some information about the interaction. Figure 5 shows the extent to which a doubling of the true interaction strength used for data generation yields a concomitant increase in the point estimates from each method tested. For both GAMs and transfer entropy, we consistently find a significant, positive relationship between the true interaction strength and the inferred point estimates, suggesting that these methods can, to some extent, characterize the relative strength of an interaction on the basis of observed case data. As above, performance was better for longer-lasting interactions. However, it is important to note that, for transfer entropy, the variation in the magnitude of point estimates across all true interaction strengths is small. For GAMs, the differences in magnitude are more substantial, although a large amount of overlap still exists (Figure S5). Furthermore, a doubling in the true interaction strength consistently leads to a much smaller increase in the point estimates returned by these methods, particularly for short-lived interactions. Thus, while these methods may correctly infer the relative interaction strength among many datasets, they are unlikely to yield meaningful conclusions about absolute interaction strength based on real-world datasets. Encouragingly, the sign of the true interaction and of the estimate returned by the GAM approach overwhelmingly agree (Figure 4, Figure S5). For Granger causality, a significant association exists for stronger interactions, but the association for shorter interactions is null. Interestingly, for both correlation coefficients and CCM, we often find a significant negative relationship, indicating that, for these methods, larger point estimates are found for data generated using weaker interactions.

**Figure 5.**
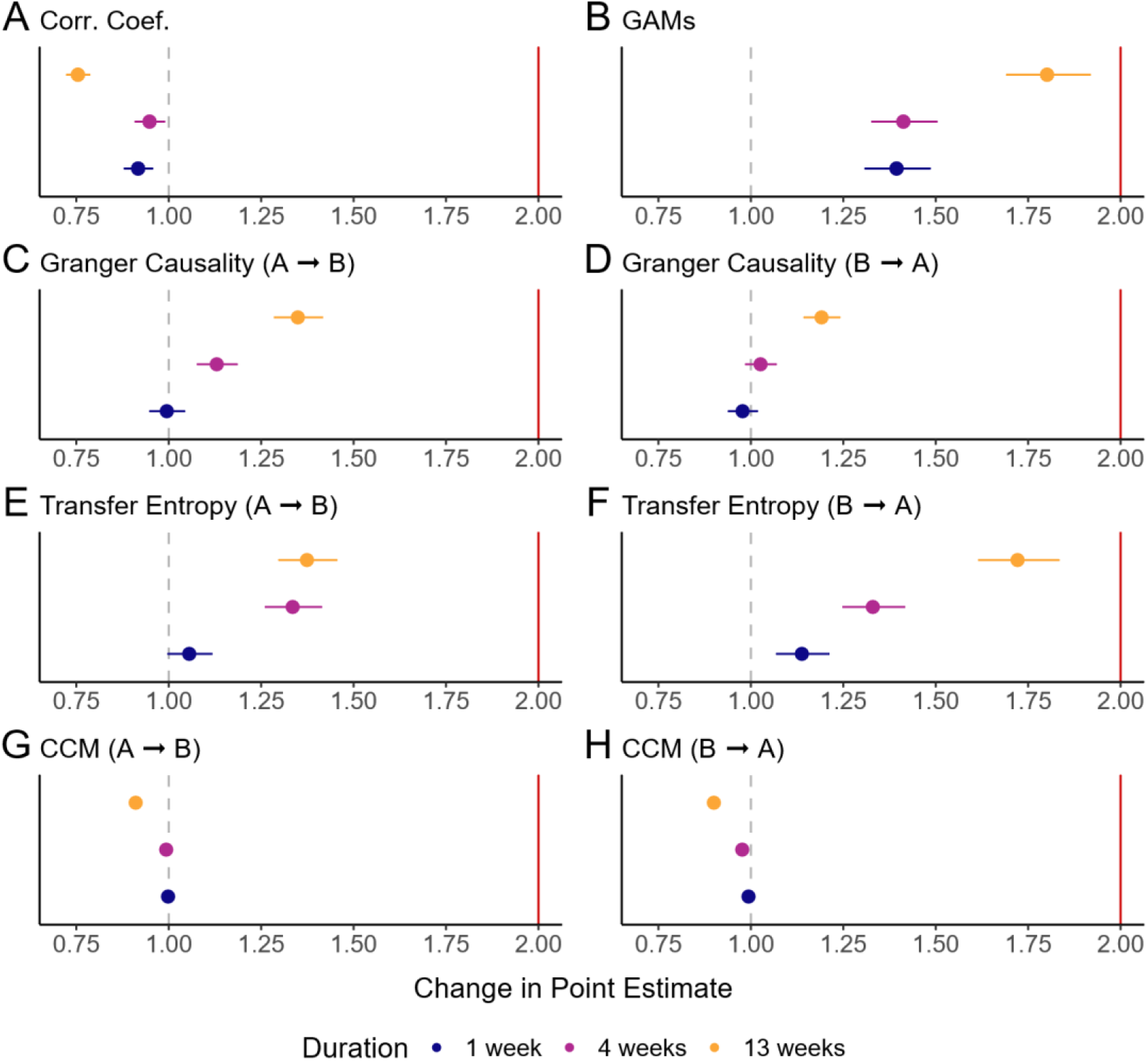
Multiplicative change in point estimates returned in response to a doubling of true interaction strength. Results are shown for all methods ((A) correlation coefficients, (B) GAMs, (C) Granger causality (impact of A on B), (D) Granger causality (impact of B on A), (E) transfer entropy (A on B) (F) transfer entropy (B on A), (G) CCM (A on B), (H) CCM (B on A)). Points show the effect estimate obtained from the linear regression model on point estimates from each method, and lines show the 95% confidence intervals. Point color indicates the true duration of the interaction. The red vertical line at 2.0 indicates the expected result if methods work perfectly (i.e., a doubling in the point estimates when the true strength is doubled), while the dotted gray vertical line at 1.0 indicates no change. Because these results are based only on the point estimates and not on whether the results are significant, results for both CCM methods are equivalent, and are therefore not shown separately.

### Methods displayed inconsistent results concerning the presence and strength of the interaction between influenza and RSV when applied to real-world data

In a previous study, we fit a mechanistic model of influenza and RSV cocirculation to surveillance data from Hong Kong and Canada, and found evidence of a negative interaction in both locations. When confronted with these same real-world data, we find that all methods tested in the current work were able to detect evidence of an interaction effect of influenza on RSV in Hong Kong. For the effect of RSV on influenza in Hong Kong, and for both interaction directions in Canada, on the other hand, conclusions were not always consistent between methods (Figure 6). As in our simulation study, GAMs performed well, identifying a significant negative interaction in both Hong Kong and Canada (Figure 6B). Both Granger causality and transfer entropy were able to identify the effect of influenza on RSV in both locations, but typically failed to identify an impact of RSV on influenza (Figure 6C-D). However, methods disagreed regarding the relative strength of these interactions across locations: Granger causality suggests a stronger interaction effect in Canada, transfer entropy suggests the opposite, and GAM results are consistent with a similar interaction strength in both Hong Kong and Canada. Finally, while CCM typically identified an interaction effect in Hong Kong, both implementations struggled to find evidence of an interaction in Canada, especially for the effect of influenza on RSV (Figure 6E).

**Figure 6.**
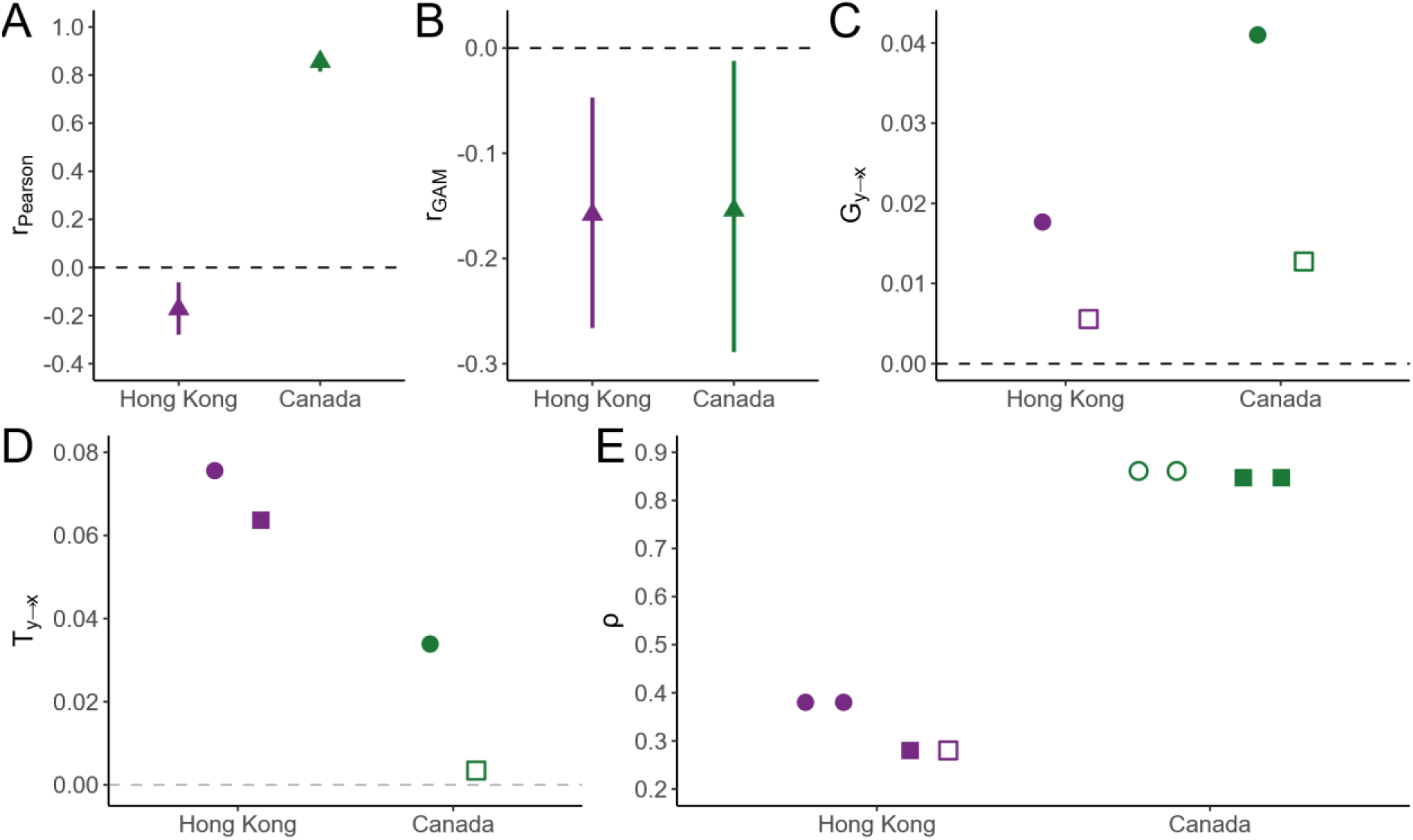
Point estimates returned by each method when confronted with influenza and RSV data from Hong Kong and Canada. Results for Pearson correlation coefficients are shown in (A), GAMs in (B), Granger causality in (C), transfer entropy in (D), and CCM in (E). Filled points represent statistically significant results (p < 0.05; for GAMs only, 95% confidence intervals not including 0), while open points represent results that were not significant. Circles show results for the interaction effect of influenza on RSV, and squares show the effect of RSV on influenza; for methods incapable of distinguishing direction (correlation coefficients and GAMs), triangles are shown. Results from Hong Kong are shown in purple, while results from Canada are shown in green. Horizontal dotted lines are plotted at values of 0. For correlation coefficients and GAMs, 95% confidence intervals are shown as vertical lines; other methods do not return confidence intervals. For CCM, results for the implementation using convergence to indicate significance are shown on the left, and results using seasonal surrogates are shown on the right; since these approaches only differ in how they evaluate significance, point estimates do not change depending on the method.

## Discussion

Although pathogen-pathogen interactions are common, they are notoriously difficult to study in human populations. Due to the nonlinear dynamics underlying infectious disease transmission, the pervasiveness of confounding, and the inherent complexity of interactions themselves, many simple statistical methods fail when applied to questions about interactions (19,20). Here, we tested five non-mechanistic statistical methods and evaluated their ability to identify and characterize the interaction between two pathogens using synthetic data. We found that even these more sophisticated methods often failed to detect whether an interaction affecting the force of infection was present at all. In cases where an interaction was present, information about the relative strength of the interaction provided by these methods was ambiguous at best.

Surprisingly, out of all of the methods tested, GAMs achieved the highest performance: in addition to correctly identifying the presence or absence of an interaction across the full range of interaction parameters tested, GAMs could also provide information about the sign and relative strength of an interaction. This was true even though this is fundamentally a regression-based approach, which is not expected to perform well in causal inference even if shared seasonality is controlled for (17). Furthermore, although we allowed the GAMs to flexibly control for the seasonality in observed case data, seasonality in observations does not necessarily correlate with seasonality in the underlying force of infection (18). It is unclear why GAMs exhibit such high performance. One possibility is that, because GAMs cannot consider the effect of A on B and of B on A separately, they are inherently observing the interaction effects of both pathogens simultaneously. This could give them an advantage in the case of a symmetric interaction. However, in a sensitivity analysis where the effect of pathogen B on pathogen A was set to 1.0 (i.e., no effect) for all simulations, we observed similar accuracy (Figure S6). Of course, the fact that GAMs are incapable of determining the direction of an interaction remains a significant weakness. Furthermore, our implementation of GAMs assumes a regular, seasonal cycle for both pathogens, and accuracy is highly dependent on the strength of the underlying seasonal forcing (Figure S7).

In contrast, although several of the tested methods (Granger causality, transfer entropy, and CCM) were developed to perform causal inference on time series data, they displayed notably worse performance. While these results may seem counterintuitive, they are in line with past work. Cobey and Baskerville (36) found that CCM often incorrectly identified cross-immunity between strains where none was modeled. When applied to real-world data on several childhood pathogens in two cities, no common interaction effects were identified between locations. Likewise, Randuineau (37) found that neither Granger causality nor transfer entropy could correctly characterize the direction or strength of unidirectional interactions. It is interesting to note that, despite being theoretically better-suited to data from systems with underlying deterministic dynamics (24), CCM performs similarly to Granger causality overall, and is slightly outperformed by transfer entropy. This echoes the results of Barraquand et al. (30), who found that Granger causality and CCM were often in agreement when identifying interactions in ecological data. Results were largely unchanged when we considered 20 (rather than 10) years of data (Figure S8), perhaps because, due to the cyclical nature of our data, little new information is gained by examining additional seasons. In the case of CCM using seasonal surrogates, the inclusion of an additional decade of data made it almost impossible to identify null interactions (Figure S8).

We observed similar results when methods were applied to real-world data. Here again, GAMs identified a significant, negative interaction between influenza and RSV in both locations, while other methods were less able to consistently detect relationships between the two viruses. Methods were generally more likely to detect an effect of influenza on RSV than of RSV on influenza. Interestingly, CCM, which is arguably the method best rooted in causal theory out of those we tested, struggled the most: neither implementation was able to identify the interaction effect of influenza on RSV in Canada, while both Granger causality and transfer entropy could.

We emphasize that our results do not imply that these methods are inherently ineffective, and our work is not intended to devalue them. Indeed, these methods are successfully applied in several other contexts (24,29,30,35,108). Rather, we find that these methods perform poorly in the specific context of pathogen-pathogen interactions, and it is important to consider why this is the case. The existence of shared seasonal forcing between our two pathogens is likely a key factor. Past work has shown that both Granger causality and CCM have a high false-positive rate when a strongly autocorrelated shared driver is at play, even when the shared driver is controlled for (30). We tested two implementations of Granger causality and transfer entropy: one accounting for the strength of the shared driver and one ignoring it. While controlling for the strength of seasonal forcing greatly reduced the number of false positives identified by both Granger causality and transfer entropy (Figure S3), specificity remained low, particularly for tests of the effect of pathogen B on pathogen A, suggesting that fully accounting for the influence of a shared driver is challenging. CCM theoretically accounts for confounding implicitly (24). However, as mentioned above, CCM performed no better than Granger causality or transfer entropy. Notably, our sensitivity analyses revealed no consistent relationship between the extent of shared forcing and method performance (Figure S7), suggesting that even very slight confounding can be a substantial problem.

This is concerning, as shared seasonal drivers are incredibly common among infectious diseases, including among those hypothesized to interact (109,110). For example, evidence suggests that transmission of both influenza and RSV, on which the pathogens in our model are based, is higher when temperatures are lower, and when humidity is either low or very high (60–62,64). Temperature and humidity are also likely to play a role in the transmission of SARS-CoV-2 (111), as well as seasonal coronaviruses (112) and rhinoviruses (113). Because this is a simulation study, we were able to control for the exact strength of seasonal forcing over time. However, in real-world outbreaks, the exact relationship between potential drivers (e.g., temperature) and pathogen transmission rates is not known, and, as explained above, seasonal patterns in forcing cannot necessarily be inferred from seasonality in pathogen incidence (17,18). We had reasonable success applying Granger causality and transfer entropy to real-world influenza and RSV data while controlling for absolute humidity, at least when it came to detecting the effect of influenza and RSV. However, RSV and especially influenza are two pathogens for which the underlying climatic drivers are relatively well-studied, which is not the case for all pathogens.

The role of confounding in general should also be discussed. Both Granger causality and transfer entropy, as well as regression-based methods like GAMs, assume that all relevant confounders are observed and accounted for (17,91,95). We are able to control for climate forcing, as weather data are widely collected and often freely available. However, we are unable to control for the proportion of the population susceptible to infection with each pathogen, a quantity that has a critical impact on transmission dynamics, and which is typically unobserved in reality (17,38). Furthermore, while we can be sure that weather is the only extrinsic variable driving transmission in our simulation study, this is not the case in real life. For example, given the apparent prevalence of pathogen-pathogen interactions, failure to account for other pathogens that interact with both pathogens of interest could lead to invalid results. Unlike Granger causality and transfer entropy, where confounding variables must be explicitly defined, GAMs control for patterns in the data resulting from confounding, while remaining mostly naïve to the nature of these confounders. While it could be argued that this gives GAMs an advantage, and may partially explain their success here, we emphasize again that there is no guarantee that patterns in observed data will be correlated with patterns in the underlying forcing functions (18). Theoretically, approaches that control for confounding directly are expected to yield more accurate results.

The impact of observation noise on method performance should also be considered, as the utility of these methods may be limited by data that are not perfectly observed (24,91). Unfortunately, for many infectious diseases, real-world outbreak data are notoriously error-laden: surveillance systems relying on healthcare utilization and testing will miss mild and asymptomatic cases, and sampling effort is often much lower outside of the outbreak season (3). Interestingly, our sensitivity analyses suggest that reducing observation noise to zero does not have a purely beneficial impact on method performance. Rather, lower observation noise tended to reduce the number of false negatives, but increased false positives, while increasing observation noise had the opposite effect (Figure S8). Furthermore, this pattern was not consistent across all methods, suggesting that the exact impact of noisy observations is difficult to predict. In contrast, modifying the amount of process noise had little effect on performance (Figure S8), a result that is mostly in line with Cobey and Baskerville (36).

It is also worth discussing that, although all of the interactions we modeled were symmetric, most methods were more likely to correctly classify the effect of pathogen A on pathogen B than the effect of B on A. This result held for both simulated and real-world data. This could be, in part, due to the seasonal driver. CCM is known to be unreliable when a downstream variable becomes synchronized to a forcing variable (24); in cases involving a shared driver, this is because this leads to data (here, observed cases data) that closely resemble the driver, potentially leading to false positives (36). In our simulated data, the strength of seasonal forcing is much more highly correlated with observed cases of pathogen B than of pathogen A, perhaps explaining the discrepancy in our results. This was not the case in our real-world data, where RSV outbreaks were only slightly but not significantly more strongly correlated with absolute humidity data. However, we note that influenza outbreaks peaked before RSV outbreaks in almost every season contained in our observed data, which could have made finding an effect of RSV on influenza more difficult. Again, we emphasize that this issue is not unique to the methods tested here, and that we experienced similar difficulties when fitting a mechanistic model to these data (3). Requiring the optimal cross-map lag to be negative as an additional significance criterion can mitigate this issue (36,104), but, for systems like seasonal infectious disease outbreaks, which have high periodicity, this approach is not valid (114). Because most pathogen-pathogen interactions are not well-understood, it is often unclear whether the interaction is uni- or bidirectional. For this reason, failure to accurately assess directionality represents a significant drawback of these methods. Even GAMs, which performed very well otherwise, cannot help here, as they offer no information about directionality.

Ultimately, the inherent complexity of pathogen-pathogen interactions may explain why many of the tested methods perform poorly here. Methods like Granger causality, transfer entropy, and CCM were originally developed with direct interactions between individual state variables in mind. In our transmission model, however, the interaction effect depends simultaneously on several model states: the number of total new infections with one pathogen is modulated by the total number of current and recent infections with the other pathogen. The issue is potentially amplified by applying the methods to outputs from the observation model, which aggregates over several states and returns underreported and error-laden case counts. On a similar note, these observed data are related to the interaction of interest through a long and indirect causal chain (see Vignette 1 in (17) for a related discussion): individual-level processes impact how likely one is to become infected with or to transmit a second pathogen (modeled here as a change in the force of infection), which in turn will impact the total number of people in a population who become infected, which is itself observed with noise. For these reasons, it is perhaps unsurprising that many of the tested methods struggle to infer pathogen-pathogen interactions. Indeed, methods like Granger causality and CCM perform well for simpler, more direct processes like, for example, predator-prey interactions (24,30). That said, these methods are already being applied to infer interactions from real-world data (39,40). We therefore reemphasize our point from the Introduction that studies explicitly evaluating these methods in this context remains an urgent necessity.

Given these challenges, none of the tested methods appear to be promising tools for characterizing pathogen-pathogen interactions on the basis of infectious disease outbreak data. Approaches utilizing mechanistic models of infectious disease transmission, like the one used to generate the synthetic data for this study, may represent an attractive alternative. By using equations to describe the transmission of pathogens throughout a population, these models explicitly account for nonlinear dynamics, as well as for potential confounders, including unobserved state variables like the proportion susceptible. Models can then be fit to surveillance data to gain insight about mechanisms of interest, including interaction effects. Although this can be computationally intensive, and improvements in the efficiency of methods for model fitting are necessary to handle, for example, models of greater than two cocirculating pathogens (10), mechanistic modeling methods show clear promise. In fact, this approach has already been used to better understand interactions between influenza and RSV (3,115), influenza and *S. pneumoniae* (13,116), influenza and SARS-CoV-2 (8), RSV and both hMPV and human parainfluenza virus (HPIV) (10,117), and different serotypes of dengue (118) and cholera (71). A key advantage of these methods over the ones tested in this work is that they can provide quantitative estimates and confidence intervals for specific characteristics of the interaction, such as its strength and duration.

We acknowledge several limitations of the current work. In particular, we have focused here on interactions between two epidemic viruses with short infectious periods and with strong periodicity in their transmission dynamics. The methods tested here may perform better for, for example, interactions involving endemic colonizing bacteria, for which carriage may persist for several weeks and prevalence is consistently high (1,2). Likewise, a modeling study found that year-to-year variability in peak incidence was critical in allowing interaction effects to be inferred from population-level data (13); more promising results may therefore be expected for pathogens showing less regular outbreak patterns. Our model was also more likely to produce outbreaks where pathogen B peaked before pathogen A (81% of datasets where no interaction was modeled) than the other way around, which could have impacted our ability to infer in particular the impact of pathogen A on pathogen B (3). However, we found little influence of the difference in peak timing between pathogens on the ability of the tested methods to identify the impact of pathogen A on pathogen B (Figure S9). Only Granger causality performed better for datasets where pathogen A peaked first, and the effect was slight; interestingly, CCM using seasonal surrogates performed slightly worse for datasets where pathogen A peaked first. Finally, although we have tested several popular methods, there are others that we have not considered (119,120). Our results do not necessarily imply that there are no non-mechanistic methods available that could provide accurate information about interactions. Rather, they emphasize the importance of rigorously testing any existing or novel methods, using an approach similar to that described in this work, before these methods are used to draw conclusions about pathogen-pathogen interactions from real-world data.

We have shown that, when confronted with infectious disease outbreak data, even sophisticated nonmechanistic methods developed with causal inference in mind are incapable of reliably identifying and characterizing pathogen-pathogen interactions. We conclude that these methods are not appropriate tools for understanding interactions on the basis of outbreak data, at least for interactions between highly seasonal, acute viruses like the one modeled here, and that interaction effects inferred using these methods should be viewed with an abundance of caution. Rather, a combination of laboratory studies and mechanistic modeling approaches is likely to be necessary to fully characterize both the individual- and population-level impact of pathogen-pathogen interactions.

## Acknowledgments

The authors gratefully acknowledge Anabelle Wong, Pietro Gemo, and Franziska Frederking for their helpful comments on the manuscript.

## Supporting Information

**S1 Text. Supplementary methods and results**.

