## Supplementary material for "The limitations of non-mechanistic methods for characterizing pathogen-pathogen interactions: A simulation study": S1 Text

### S1 Text. Supplementary Materials

#### 1. Model Development and Choice of Parameter Values

There is little information in the literature concerning overdispersion in reporting or the extent of superspreading for RSV in particular. However, as RSV outbreaks are typically more regular and less noisy than outbreaks of influenza, we simply halved the values for $k_A$ and $\beta_{sd\_A}$ to obtain $k_B$ and $\beta_{sd\_B}$, respectively.

In our model, previously infected individuals are able to lose their immunity over time and return to the susceptible compartments, allowing reinfection. However, due to the way interactions are modeled, recovered individuals must pass through both the T compartments (where they are recovered but still subject to interaction effects) and the R compartments (where the interaction no longer plays a role) before they are susceptible again. For this reason, the total time until loss of immunity is determined not only by $\omega$, but also by the duration of the interaction ($\delta$). To correct for this, we adjust the rate of loss of immunity within the model for each pathogen such that:

$$\omega' = 1 / (\frac{1}{\omega} - \frac{1}{\delta})$$

This correction ensures that the time required to pass through both the T and R compartments will be, on average, the desired duration of immunity, as listed in Table 1.

As described in the main text, surges in immunity loss for pathogen A were modeled as occurring once per year, with the exact timing drawn from a normal distribution with mean 12 and standard deviation 4, such that surges occur on average 12 weeks into the outbreak (i.e., during the thirteenth week). Because we are working with purely synthetic data, the exact choice of surge timing is not critical. However, Yaari et al. report that these surges tend to precede the initial rise in epidemic activity each year (1). Thus, it makes sense to place the surges several weeks prior to the peak in transmissibility (here, $\phi$ = 26), which tends to occur shortly before peak incidence. The size of surges ranged from 5-30% of the recovered population, as described in the main text, and was drawn from a uniform distribution. Values of $R_{effA}$, $R_{effB}$, $\omega_B$ for each of the 100 parameter sets were chosen from the ranges in main text Table 1 using Latin hypercube sampling.

#### 2. Additional Methodological Details for Statistical Inference

Although the data-driven, non-mechanistic methods tested here require fewer assumptions about the system under study than more mechanistic approaches, some knowledge is still necessary for parameterization. For Granger causality, for example, an appropriate order for the VAR models must be selected. For transfer entropy, users must specify the history length for both the predictor and outcome variables. For CCM, the embedding dimensions of both the predictor and outcome variables are required. Although some of these parameters can be chosen automatically by selecting the values that maximize within-pathogen predictive ability, users must still provide sensible upper bounds.

For our particular application to infectious disease data, we are primarily interested in a) how long can a past case of pathogen A (or pathogen B) impact how many cases of the same pathogen we observe during the current week *t*, and b) for how long can a past case of pathogen A (or pathogen B) impact how many cases of the potential interacting pathogen we observe during the current week *t*?

To answer the former question, we look at generation times, defined as the time between infection of a primary case and subsequent infection of a secondary case (2). If generation time for a pathogen is, for example, 3 weeks, this suggests that current case counts can be affected by cases that occurred up to three weeks previously. The distribution of the generation time for a pathogen, then, will depend on both the latent period, which all newly-infected cases must pass through before becoming infectious, and the infectious period, the period of time during which onward transmission is possible. For a given primary and secondary case of pathogen A, for example, the generation period can be calculated as:

$$T_E + UT_I$$

where:

$$T_E \sim Exp(\sigma_A)$$
$$T_I \sim Exp(\gamma_A)$$
$$U \sim U(0, 1)$$

Here, $T_E$ and $T_I$ represent random draws from the distribution of possible latent and infectious periods, respectively. Meanwhile, $U$ accounts for the fact that a secondary infection can occur at any point during the primary case's infectious period.

By generating a large number of possible generation times using the equations above, we find that the vast majority (>98%) of secondary cases of pathogen A will occur within 2 weeks of primary case infection; for pathogen B, this value was closer to 5 weeks. These values were used as the history lengths for each pathogen when performing transfer entropy, specifically when the pathogen in question acted as the outcome of interest. Values of 2 and 5 were also set as the maximum embedding dimensions for each pathogen when performing CCM.

In the case of a pathogen-pathogen interaction, the situation is more complex. A past infection with pathogen A will not only influence subsequent cases of pathogen B for the period of time over which the case of pathogen A is transmitting, but also potentially for several weeks after recovery, depending on the timeframe of the interaction. For this reason, we chose to add together the generation times of both pathogens (2 weeks and 5 weeks), as well as the maximum possible duration of the interaction (13 weeks), to yield 20 weeks. This value was taken to be the maximum order of the VAR models used in the Granger causality analyses, the history length for predictor variables when performing transfer entropy, and the maximum lag when running CCM.

Finally, when generating seasonal surrogates to evaluate the statistical significance of CCM results, users must specify $\alpha$, the standard deviation of the normally-distributed noise to be added to the surrogate data. Before running our final analyses, we tested several values of $\alpha$ to determine their influence on CCM accuracy. We found that, as $\alpha$ increases, sensitivity also increases, but specificity decreases (results not shown). Since most of our methods struggle much more with returning false positives than false negatives, we prioritized maximizing specificity and chose to set $\alpha$ to 0 for both pathogens.

**3. Sensitivity Analyses**

In general, sensitivity analyses involved varying several key model parameters, one at a time, and generating new synthetic data. More specifically, we performed sensitivity analyses to evaluate the impact of:

- Asymmetric interactions ($\theta_{\lambda_B}$= 1.0; GAMs only)
- Lower process noise ($\beta_{sd\_A}$ = 0.01, $\beta_{sd\_B}$ = 0.005)
- Higher observational noise ($k_A$ = 0.1; $k_B$ = 0.05)
- No observational noise ($k_A$ = $k_B$ = 0)
- Longer time series available for inference (20 years of weekly data)
- Varying amplitude of seasonal forcing ($\alpha$ = 0.05, 0.3, 0.4, 0.5)

All statistical methods were then applied to these data as described in the main text. When the amplitude of seasonal forcing was low ($\alpha$ = 0.05), more of the resulting data was not stationary (n = 24, vs. n = 2 in the main analysis), and GAMs were less likely to achieve a good-quality fit (13.9% of datasets removed, vs. 2.1% in the main analysis). Data were also less likely to be stationary for $\alpha$ = 0.4 or 0.5 (n = 12). For all other analyses, stationarity and model fit appeared similar to those achieved in the main analysis. Results from these analyses are discussed throughout the main text.

As described above, we initially chose the upper bounds for several method parameters based on the generation times of influenza and RSV. However, since future cases of a pathogen will depend on both the number of infected and the number of susceptible individuals in the population, and the duration of immunity for our modeled pathogens ranges from 7.2 to 12 months, past cases of a given pathogen may influence cases at time *t* for significantly longer than the generation time. We therefore repeated our analyses using a value of 37 weeks (the median time to loss of immunity given an exponential distribution with rate = $1/\omega$, or 1/52.25). Specifically, this value was used as the history length of the outcome variable (transfer entropy) and the maximum embedding dimension (CCM) for both pathogens; a value of 50 (37 + 13 weeks maximum interaction duration) was used for the maximum order of VAR models (Granger causality), the history length of the predictor variable (transfer entropy), and the maximum lag (CCM).

These changes had little influence on the overall sensitivity and specificity of the tested methods, particularly for Granger causality and transfer entropy (Figure S4). For CCM, allowing higher values for the embedding dimensions and lags led to an increase in sensitivity, but a roughly equivalently-sized loss in specificity. In fact, when using convergence to assess significance, these larger parameter values led to near perfect sensitivity, but specificity of near 0. The only scenario where CCM showed clear overall improvement was for the effect of pathogen B on pathogen A when using seasonal surrogates to evaluate significance: here, sensitivity increased, while specificity remained roughly the same. However, this specificity was still incredibly low (0.197).

As detailed in the main text, GAMs that either resulted in divergent transitions or did not converge are removed from the analysis. The corresponding datasets, however, are not removed when analyzing any of the other methods we tested. To ensure that this decision did not provide GAMs with an unfair advantage, we re-analyzed our results, this time removing any datasets that resulted in poorly-fitting GAMs from all analyses. Encouragingly, this yielded no noticeable change in our results: the greatest observed change in sensitivity or specificity for the other methods relative to the main text results was 0.0053 (results not shown).

#### 4. Impact of Phase Difference on Accuracy

In simulations where no interaction was modeled, our model was much more likely to produce outbreaks with an earlier peak timing of pathogen B than pathogen A. To assess whether this impacted the tested methods' ability to correctly infer the interaction effect of pathogen A on pathogen B, we first calculated the peak timing for both pathogens for all ten years of simulated data in each of our 100 parameter sets. To obtain the average phase difference for each dataset, we subtracted the peak timing of pathogen B from the peak timing of pathogen A, such that positive values indicate an earlier peak for pathogen B, and took the median over all 10 seasons. Specifically, phase differences were calculated only for those simulations where no interaction effect was modeled. This was done to avoid any confounding by the true strength and duration of the interaction, as interactions can influence the timing of outbreaks of both pathogens (Figure 2 and Figure S2). Finally, we used GAMs to fit logistic regression models for each method. Here, the dependent variable was whether or not the interaction effect was correctly estimated for each simulated dataset, while the independent variables were 1) a smooth on the phase difference calculated when no interaction occurred (as described above) for a given parameter set, and 2) a random effect on parameter set (1-100). Results can be seen in Figure S8, and are discussed in the main text.

Supplementary Equations

$$\frac{dX_{SS}}{dt} = \omega'_A(t)X_{RS} + \omega'_B X_{SR} + \mu N - (\lambda_A + \lambda_B + \nu)X_{SS}$$

$$\frac{dX_{ES}}{dt} = \lambda_A X_{SS} + \omega'_B X_{ER} - (\sigma_A + \lambda_B + \nu)X_{ES}$$

$$\frac{dX_{IS}}{dt} = \sigma_A X_{ES} + \omega'_B X_{IR} - \left(\gamma_A + \theta_{\lambda_A}\lambda_B + \nu\right)X_{IS}$$

$$\frac{dX_{TS}}{dt} = \gamma_A X_{IS} + \omega'_B X_{TR} - \left(\delta_A + \theta_{\lambda_A}\lambda_B + \nu\right)X_{TS}$$

$$\frac{dX_{RS}}{dt} = \delta_A X_{TS} + \omega'_B X_{RR} - (\omega'_A(t) - \lambda_B + \nu)X_{RS}$$

$$\frac{dX_{SE}}{dt} = \omega'_A(t)X_{RE} + \lambda_B X_{SS} - (\lambda_A + \sigma_B + \nu)X_{SE}$$

$$\frac{dX_{EE}}{dt} = \lambda_A X_{SE} + \lambda_B X_{ES} - (\sigma_A + \sigma_B + \nu)X_{EE}$$

$$\frac{dX_{IE}}{dt} = \sigma_A X_{EE} + \theta_{\lambda_A}\lambda_B X_{IS} - (\gamma_A + \sigma_B + \nu)X_{IE}$$

$$\frac{dX_{TE}}{dt} = \gamma_A X_{IE} + \theta_{\lambda_A}\lambda_B X_{TS} - (\delta_A + \sigma_B + \nu)X_{TE}$$

$$\frac{dX_{RE}}{dt} = \delta_A X_{TE} + \lambda_B X_{RS} - (\omega'_A(t) + \sigma_B + \nu)X_{RE}$$

$$\frac{dX_{SI}}{dt} = \omega'_A(t)X_{RI} + \sigma_B X_{SE} - \left(\theta_{\lambda_B}\lambda_A + \gamma_B + \nu\right)X_{SI}$$

$$\frac{dX_{EI}}{dt} = \theta_{\lambda_B}\lambda_A X_{SI} + \sigma_B X_{EE} - (\sigma_A + \gamma_B + \nu)X_{EI}$$

$$\frac{dX_{II}}{dt} = \sigma_A X_{EI} + \sigma_B X_{IE} - (\gamma_A + \gamma_B + \nu)X_{II}$$

$$\frac{dX_{TI}}{dt} = \gamma_A X_{II} + \sigma_B X_{TE} - (\delta_A + \gamma_B + \nu)X_{TI}$$

$$\frac{dX_{RI}}{dt} = \delta_A X_{TI} + \sigma_B X_{RE} - (\omega'_A(t) + \gamma_B + \nu)X_{RI}$$

$$\frac{dX_{ST}}{dt} = \omega'_A(t)X_{RT} + \gamma_B X_{SI} - \left(\theta_{\lambda_B}\lambda_A + \delta_B + \nu\right)X_{ST}$$

$$\frac{dX_{ET}}{dt} = \theta_{\lambda_B}\lambda_A X_{ST} + \gamma_B X_{EI} - (\sigma_A + \delta_B + \nu)X_{ET}$$

$$\frac{dX_{IT}}{dt} = \sigma_A X_{ET} + \gamma_B X_{II} - (\gamma_A + \delta_B + \nu)X_{IT}$$

$$\frac{dX_{TT}}{dt} = \gamma_A X_{IT} + \gamma_B X_{TI} - (\delta_A + \delta_B + \nu)X_{TT}$$

$$\frac{dX_{RT}}{dt} = \delta_A X_{TT} + \gamma_B X_{RI} - (\omega'_A(t) + \delta_B + \nu)X_{RT}$$

$$\frac{dX_{SR}}{dt} = \omega'_A(t)X_{RR} + \delta_B X_{ST} - (\lambda_A + \omega'_B + \nu)X_{SR}$$

$$\frac{dX_{ER}}{dt} = \lambda_A X_{SR} + \delta_B X_{ET} - (\sigma_A + \omega'_B + \nu)X_{ER}$$

$$\frac{dX_{IR}}{dt} = \sigma_A X_{ER} + \delta_B X_{IT} - (\gamma_A + \omega'_B + \nu)X_{IR}$$

$$\frac{dX_{TR}}{dt} = \gamma_A X_{IR} + \delta_B X_{TT} - (\delta_A + \omega'_B + \nu)X_{TR}$$

$$\frac{dX_{RR}}{dt} = \delta_A X_{TR} + \delta_B X_{RT} - (\omega'_A(t) + \omega'_B + \nu)X_{RR}$$

$$\frac{dH_1}{dt} = \gamma_1(X_{IS} + X_{IE} + X_{II} + X_{IT} + X_{IR})$$

$$\frac{dH_2}{dt} = \gamma_2(X_{SI} + X_{EI} + X_{II} + X_{TI} + X_{RI})$$

**Equation S1. Full model equations used to generate synthetic data.** The force of infection ($\lambda_i$) is calculated by multiplying $\beta_i(t)$ by the prevalence of pathogen *i* at time *t*. Here, $\omega'_i$ represents the rate of loss of immunity adjusted for the duration of the interaction (see Supplementary Text); the dependence of $\omega'_A$ on time allows for the inclusion of surges in loss of immunity at the selected timepoints.

#### Supplementary Figures

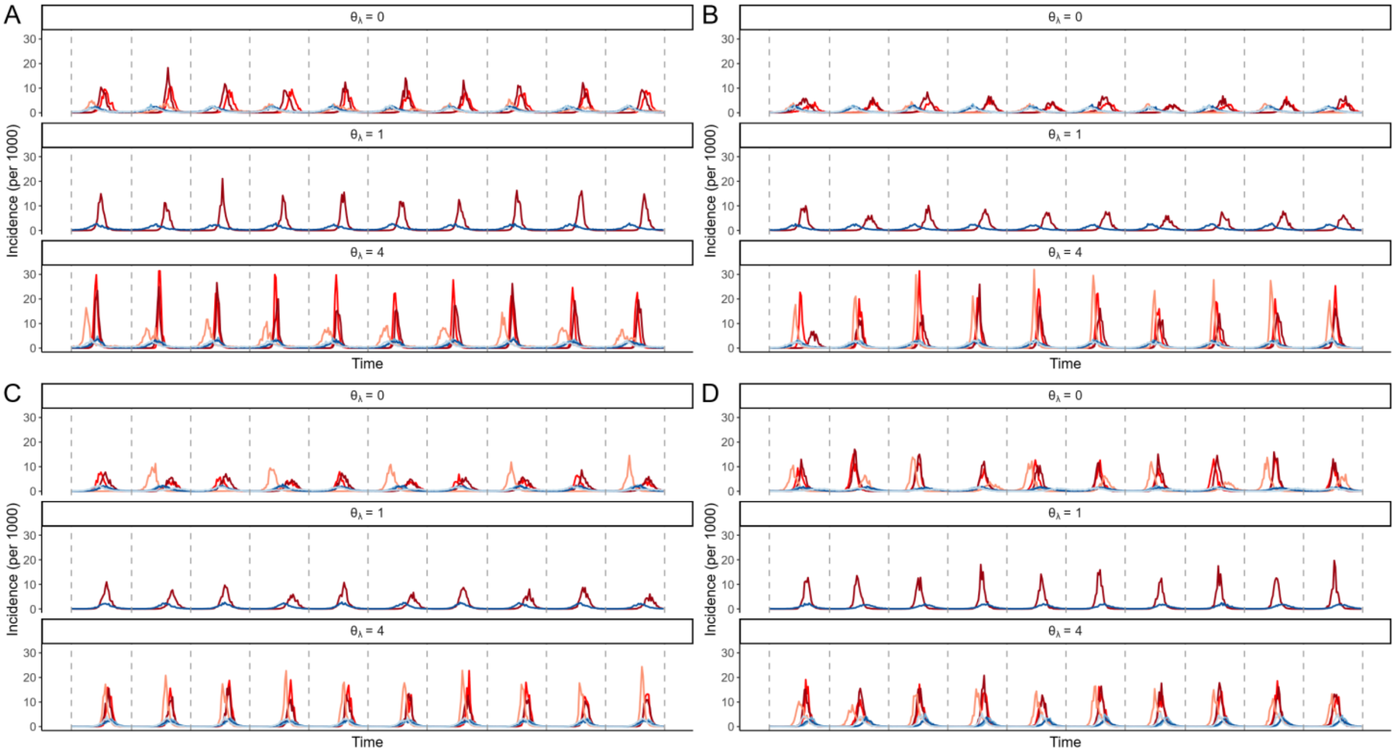


**Figure S1. Four representative data sets generated from the model.** Figure layout and colors are as in Figure 2. In each panel, interaction strength ($\theta_\lambda$) and duration ($7/\delta$) were varied, while all other parameters were held constant for that dataset. Specifically, parameter values were: (A) $Ri_1$ = 1.36, $Ri_2$ = 1.88, $1/\omega_2$ = 32.5 weeks; (B) $Ri_1$ = 1.07, $Ri_2$ = 1.95, $1/\omega_2$ = 36.8 weeks; (C) $Ri_1$ = 1.11, $Ri_2$ = 1.80, $1/\omega_2$ = 44.7 weeks; (D) $Ri_1$ = 1.38, $Ri_2$ = 1.62, $1/\omega_2$ = 45.4 weeks; all other parameters were set to the values in Table 1.

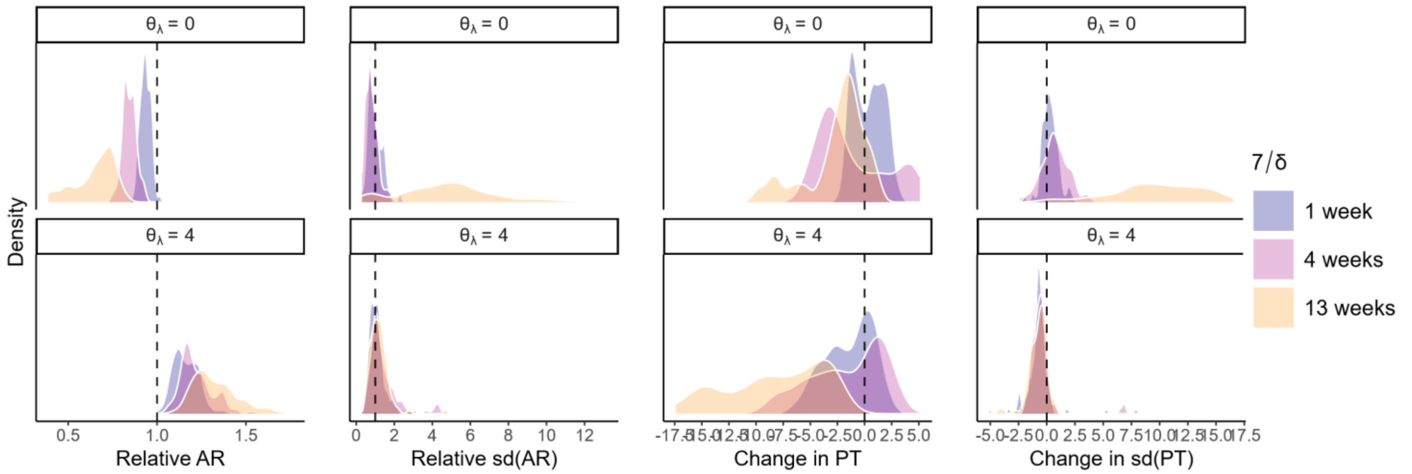


**Figure S2. Change in dynamics of pathogen A due to interaction.** Similar to Figure 2B, the distribution of the change in attack rate, year-to-year variation in attack rate, peak timing, and year-to-year variation in peak timing of pathogen A when either a strong negative (top) or strong positive (bottom) interaction is at play, relative to the dynamics when no interaction is present. Shading indicates the duration of the interaction; dotted vertical lines indicate no change.

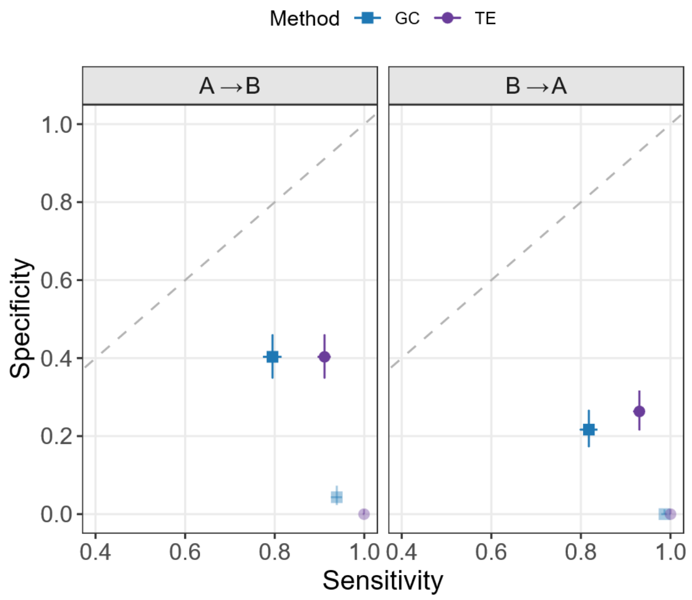


**Figure S3. Sensitivity and specificity for implementations of Granger causality and transfer entropy accounting for vs. ignoring shared seasonal forcing.** Method is indicated by both point shape and color, as in Figure 3; darker colors show the results from the main analysis, where shared seasonal forcing is accounted for, while the faded points show results from an analysis where shared seasonal forcing is not included. The crosshairs superimposed on each point represent 95% confidence intervals. As in Figure 3, the dashed diagonal lines show where sensitivity is equal to specificity. Results for the effect of pathogen A on pathogen B are shown on the left, and results for the effect of B on A are shown on the right.

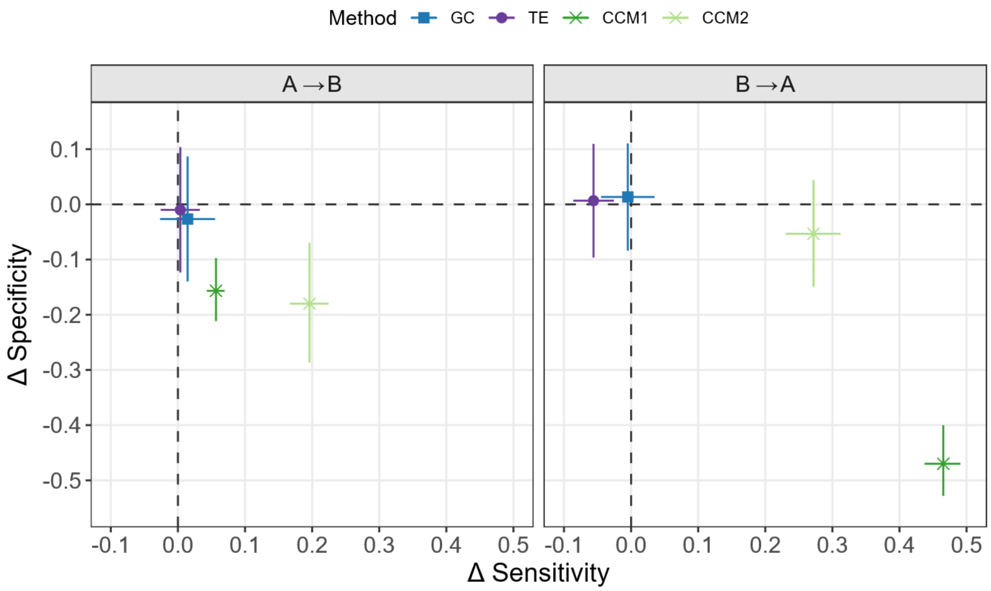


**Figure S4.** Change in sensitivity and specificity, relative to main results (Figure 3), when higher values are used as maximum orders for VAR models (Granger causality), history lengths (transfer entropy), and maximum embedding dimensions and lags (CCM). Method is indicated by both point shape and color, as in Figure 3. The crosshairs on each point represent 95% confidence intervals. The dashed vertical and horizontal lines indicate no change in sensitivity and specificity, respectively. Results for the interaction effect of pathogen A on pathogen B are shown on the left, while results for the effect of pathogen B on A are shown on the right.

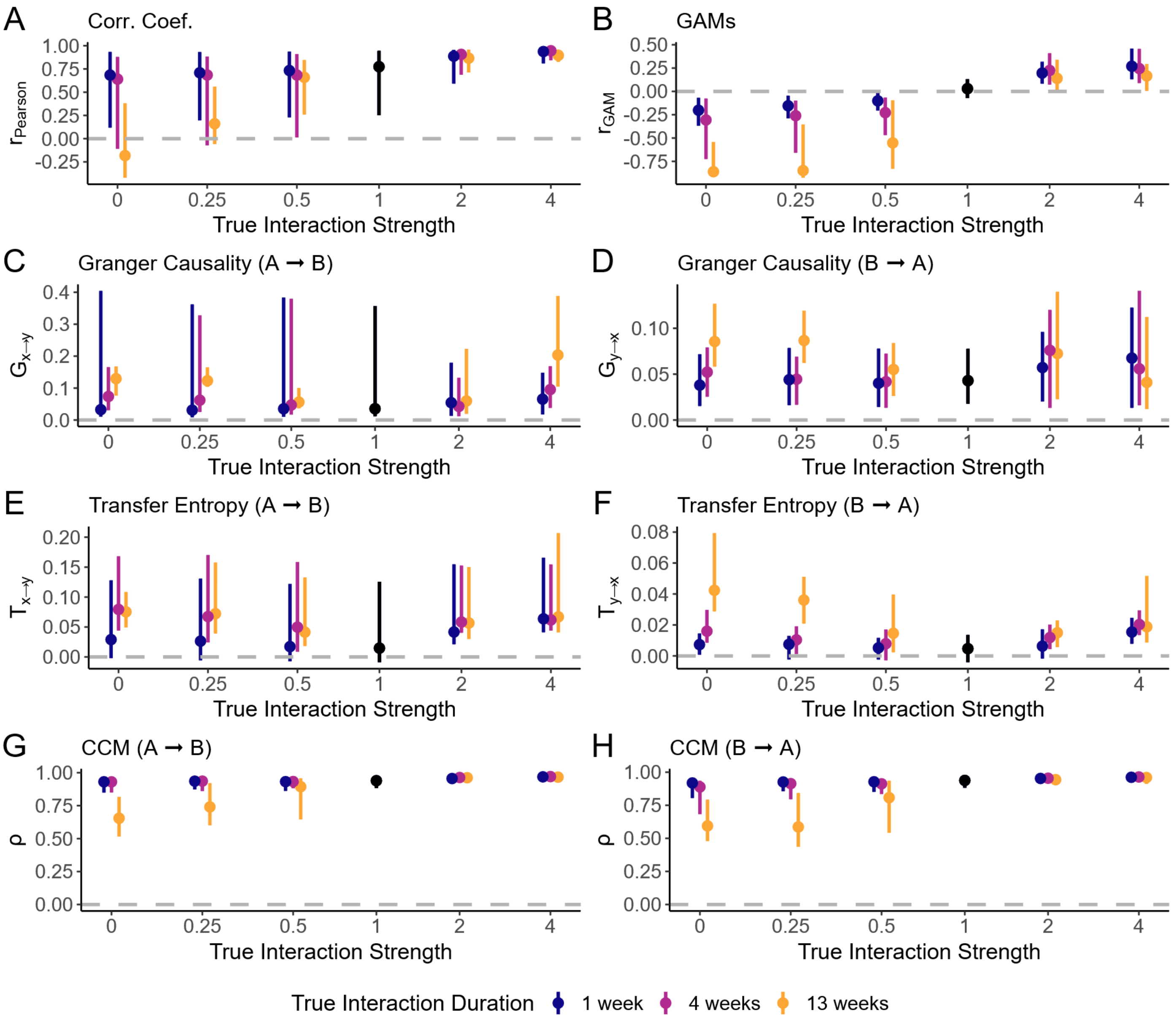


**Figure S5. Point estimates returned by each method, according to true interaction strength and duration.** Results are shown for all methods and directions (correlation coefficients (A), GAMs (B), Granger causality (C-D), transfer entropy (E-F), and CCM (G-H); results for effect of pathogen A on pathogen B shown in (C), (E), and (G), and results for effect of B on A in (D), (F), and (H)). True interaction strength is shown on the x-axis, while true direction is indicated by color. The black points and lines show results for simulations where no interaction was included; the gray dotted lines at zero indicates a null result (i.e., no interaction identified).

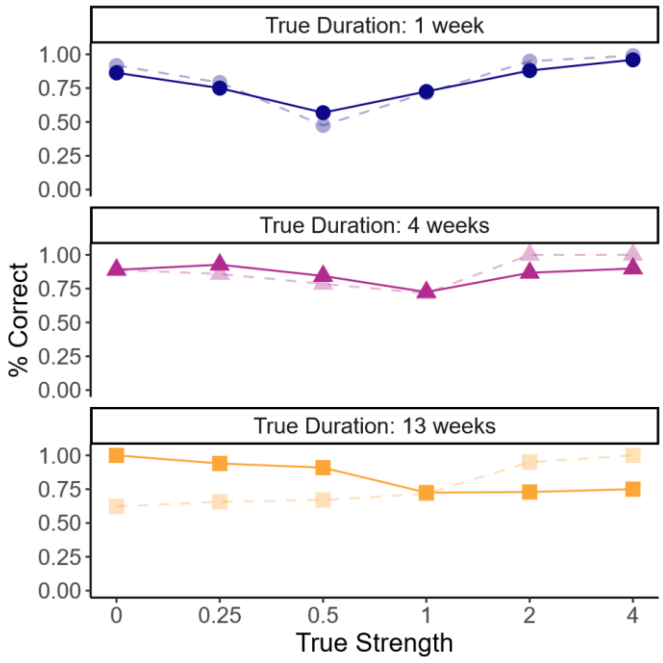


**Figure S6. Accuracy of GAMs for symmetric vs. asymmetric (unidirectional) interactions.** Results for the asymmetric interaction, where pathogen B has no effect on susceptibility to pathogen A, are shown as dotted lines; the main text results, for a symmetric interaction, are shown as solid lines. True interaction strength is shown on the x-axis, while true interaction duration is shown by panel; colors and shapes are as in main text Figure 4.

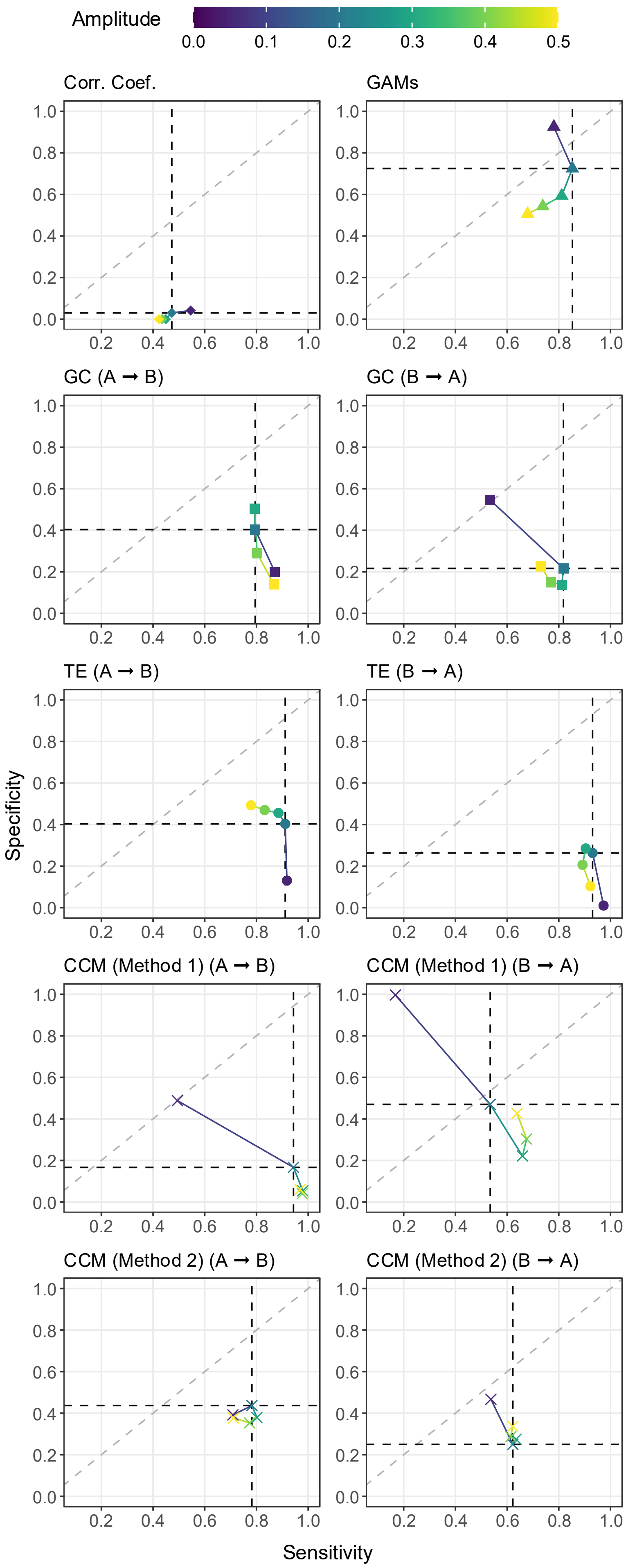


**Figure S7. Sensitivity and specificity for all tested methods as the amplitude of shared seasonal forcing is varied.** For all methods capable of distinguishing direction, the tested direction is indicated for each panel. Point color indicates the amplitude of seasonal forcing; shapes are as in Figure 3. The dotted diagonal line shows where sensitivity is equal to specificity. Dotted vertical and horizontal lines show the sensitivity and specificity, respectively, obtained in the main analysis ($\alpha$ = 0.20, also shown here by the blue point).

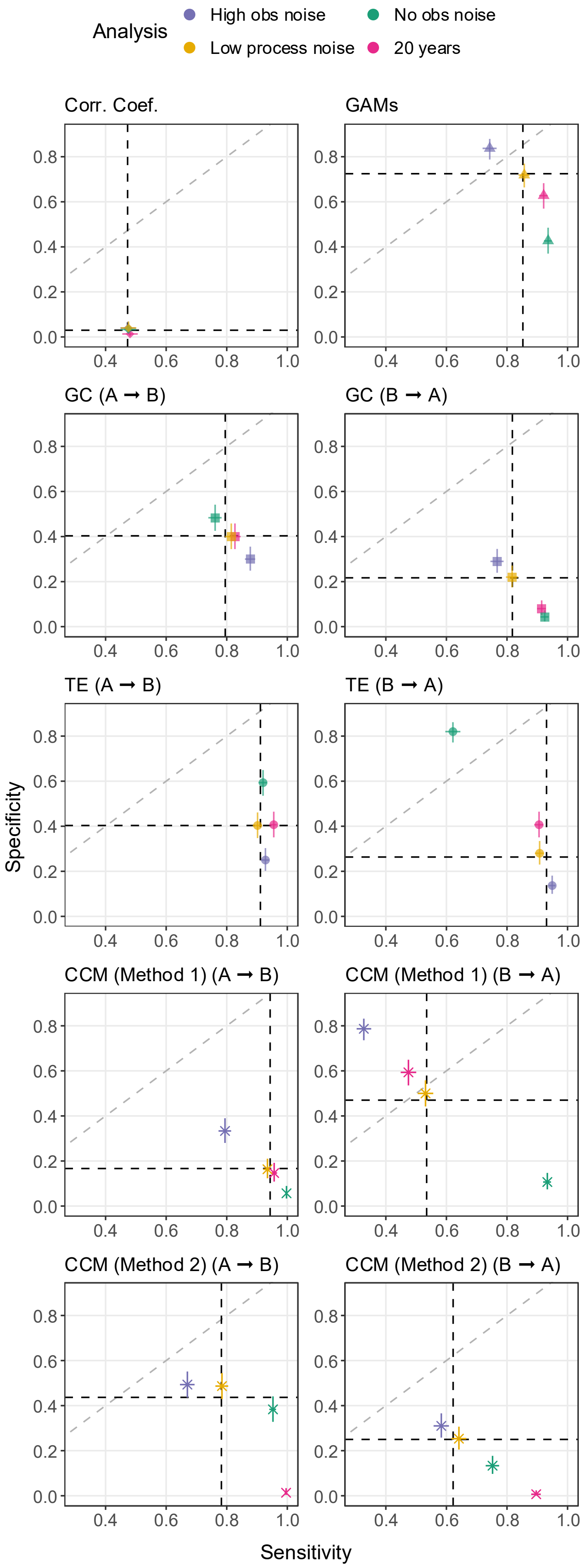


**Figure S8. Sensitivity and specificity for all tested methods for a range of sensitivity analyses varying process noise, observation noise, and the number of years of available data.** For all methods capable of distinguishing direction, the tested direction is indicated for each panel. The specific sensitivity analysis is indicated by color, shapes are as in Figure 3, and crosshairs indicate 95% confidence intervals. The dotted diagonal line shows where sensitivity is equal to specificity. Dotted vertical and horizontal lines show the sensitivity and specificity, respectively, obtained in the main analysis.

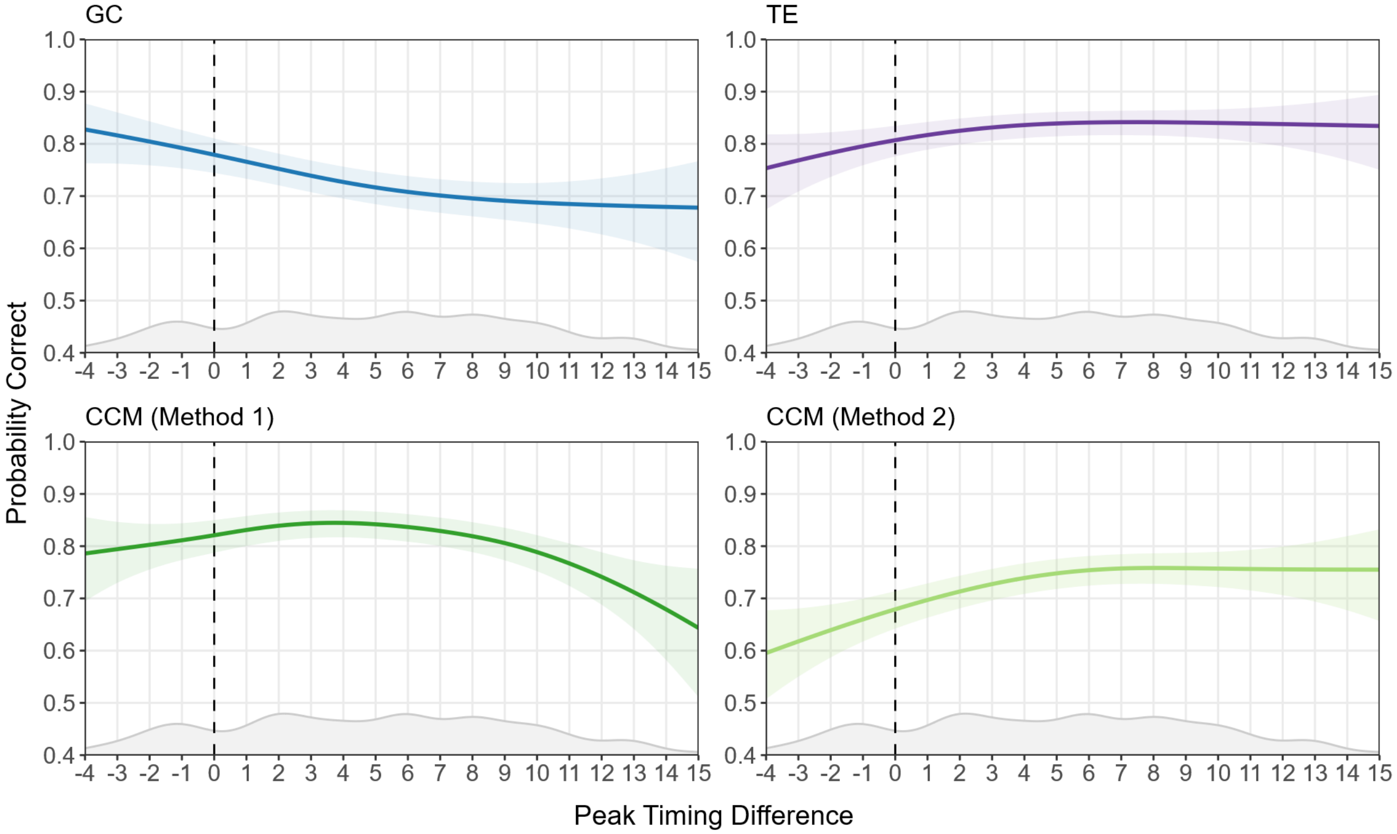


**Figure S9. Impact of the difference in peak timing between pathogens A and B on method accuracy.** Lines show the marginal effect of peak timing difference on the chance of correctly identifying the impact of pathogen A on pathogen B, for all methods capable of distinguishing directionality, after controlling for parameter set. Shaded areas represent 95% confidence intervals. Colors for each method are as in Figure 3. Positive values of peak timing difference indicate simulations where pathogen B peaked earlier in simulations with no modeled interaction, while negative values indicate simulations where pathogen A peaked first; the vertical dotted line at 0 indicates simulations where both pathogens peaked simultaneously. Density plots along the bottom of each panel show the relative frequency with which different values of peak timing difference were observed in our dataset.
